# Attraction to secreted isoamyl alcohol as a signal for beneficial commensals

**DOI:** 10.64898/2026.05.22.727293

**Authors:** J Pietropaolo, S El Khoury, L Guo, M McGee, M Shapira

## Abstract

*C. elegans,* a bacterivore living in microbially-complex environments, harbors a characteristic community of gut bacteria that contribute to its health and fitness. What determines which environmental bacteria end up as commensals is largely unknown in *C. elegans*, as in other animals. Previous work found that gut *Pantoea* isolates supported rapid worm development and infection resistance, while environmental congenerics were inferior. Notably, worms were preferentially attracted to the more beneficial gut isolates. Using bioactivity-guided fractionation and gas chromatography-mass spectrometry analysis, we identified bacterially derived isoamyl alcohol (IAA) as a secreted volatile attractant underlying this preference. Screening of worm mutants implicated AWC sensory neuron-associated genes in preferential attraction to beneficial *Pantoea* and established a causal link between IAA sensing and colonization by beneficial strains. While IAA sensing was important for initial colonization, gut-associated *Pantoea* strains ultimately outcompeted environmental congenerics over time, indicating that microbiome assembly is shaped by two complementary processes: host behavioral preference for high-IAA producers and bacterial competitive fitness within the gut. While IAA is a product of leucine metabolism and may function as a nutritional cue, we found that it could also directly enhance host infection resistance, suggesting an additional role in modulating host physiology. Finally, knockout analysis identified a bacterial branched-chain amino acid aminotransferase homolog as important for IAA production. Together, these findings identify bacterial volatile sensing as an important and underexplored mechanism shaping microbiome composition and its contributions to host fitness.

## Introduction

The gut microbiome has been a booming area of research in recent years aimed at understanding its diverse impacts on host health and fitness [1–4]. Gut microbiomes are extensive and diverse, but their composition is not random; it is key to their impact on host health and fitness, and gross deviations from the more typical composition for a host species (i.e. dysbiosis) have been causally linked to pathology, including obesity and depression [5, 6]. Further, inter-species microbiome transplants have shown to reduce host fitness [7, 8]. The shaping of a beneficial gut microbiome is affected by many factors. These include mother-to-offspring transmission, host genetics, and diet [9–16], but also environmental availability of microbes, acquired from food or through social interactions [10, 17–21]. Which factors determine environmental acquisitions is not well understood. Behavior, important in determining animal interactions with the environment, may be one such factor, but very little is known about its role in bacterial acquisitions, or the signals that may induce it. Smelling of putrefying bacteria in food deters animals from consuming it, along with potentially harmful bacteria [22, 23]. Immune mechanisms further identify and remove pathogenic bacteria [24, 25]. In contrast, hosts may be attracted to beneficial microbes, which as commensals, could increase their fitness [26]. However, very little is known about this type of bacterial acquisition or the mechanisms that underlie them. Bacterial diversity is immense; genera can include thousands of different species, and species may include thousands of strains, which are genetically different and may differ dramatically in their associated functions [27]. To what extent can animals distinguish between useful and less useful non-pathogenic species, which bacterial features are being recognized, and how?

The nematode *Caenorhabditis elegans* is seeing a growing interest as a model for microbiome research [28–34]. In addition to its advantages as a genetic model, it provides the opportunity to work with clonal populations—age-synchronized and gnotobiotic when so desired—to dissect microbiome composition and the mechanisms that shape it. *C. elegans* is a bacterivorous organism that reproduces on rotting organic matter, where it is exposed to a large diversity of bacteria [35, 36]. However, its gut microbiome was found to be distinct from the environmental diversity and presented a characteristic composition that was similar in worms raised in different environments [37–39]. Immune mechanisms play an important part in shaping the worm gut microbiomes [31, 40, 41], but *C. elegans* is also known to effectively sense and respond to a large variety of metabolites. Much of the research on sensing in *C. elegans* has focused on studying neuronal circuits and neuronal function, without consideration of the biological context of the sensed metabolites. This, however, subsequently provided mechanistic basis for understanding avoidance of pathogenic bacteria, shown in several cases to be based on sensing characteristic metabolites [42–44]. More recently, research has also shown that worms can sense and be attracted to bacterial metabolites signifying sources of nutrients or bacteria to become gut commensals [45–47].

Here, we identify a volatile metabolite secreted by gut commensals of the genus *Pantoea* that attracts *C. elegans* worms and promotes their colonization over low-secreting environmental congenerics. Members of the genus *Pantoea* were isolated from worms both in Europe and in the US [48–50]. *Pantoea* species are common in diverse environments, where they can be free-living, associated with plants, or with animals, including as endosymbionts [51–53]. Previous work in our lab isolated distinct species from the gut of worms raised in compost microcosms or from the compost environment. Gut isolates were beneficial for worm growth and infection resistance, whereas environmental isolates were less beneficial. Furthermore, worms were preferentially attracted to the beneficial gut isolates, suggesting that they could distinguish among strains based on cues linked to host benefit. Here, supported by the isolation of additional environmental strains, we identify isoamyl alcohol as the bacterial metabolite underlying this attraction, promoting colonization by high-producing commensals.

## Materials and Methods

### Worm Strains and Growth Conditions

Worms of the Bristol N2 wildtype strain were used in most experiments. Mutant strains tested for impaired sensing are described in Supplementary Materials (Table S1).

Worms of all strains were raised at 20°C on nematode growth medium (NGM), with a lawn of *E. coli* strain OP50 as food. All worm strains, as well as the *E. coli* OP50 bacteria, used as standard food, were obtained from the *Caenorhabditis* Genome Center (CGC).

Synchronized worm populations were obtained by bleaching gravid worms to release eggs and then hatching them in M9 salt solution without food, giving rise to arrested L1 larvae. Development was then allowed to progress by shifting larvae to plates with bacterial lawns [54].

### Pantoea Strains and Bacterial Growth

BIGb0393 (GenBank: GCA_037479095.1), a *P. nemavictus* strain, is one of 12 bacterial strains of the CeMbio community of worm commensals [48]; *P. cypripedii* strains V8 (GenBank: GCA_036663555.2) and T16 (GenBank: GCA_036663515.1) were previously isolated from the gut of adult worms raised in compost microcosms [50]. *Pantoea* strains T14, T15, T19, and T30 were isolated from the worms’ compost environment in the same experiments as V8 and T16. T14 (GenBank: GCF_036663535.1) was previously described [50], and identified here as *Pantoea phytostimulans*. T15, T19 and T30 (Accession Numbers: JBXGSM000000000, JBXGSN000000000, JBXGSO000000000) were characterized as part of the current study and were also identified as *Pantoea phytostimulans*. *Pantoea* strains were grown at 28°C in LB liquid media without antibiotics and with shaking. Cultures were inoculated from a saturated overnight culture (15-20% v/v inoculum) and grown for additional 18-24 hours (resulting in mid to late-stationary phase) before adding to plates or measuring metabolite concentrations.

### Whole Genome Sequencing

Single colonies of the respective strains were grown overnight in LB, and genomic DNA was extracted using a Zymo DNA extraction kit. Purified DNA was submitted to the UC Berkeley QB3 facility for whole genome sequencing on a NovaSeq X Plus. Raw reads were assembled *de novo* into contigs using SPAdes [55]. Assembly of contigs used only those larger than 1000bp, giving rise to draft genomes, which were deposited to NCBI (Table S2). Draft genomes were annotated with PROKKA [56], compared to each other with average nucleotide identity (ANI) [57] (Table S3) and MUMmer, used to align genomes and identify genetic variation [58] (Table S4). These comparisons demonstrated high similarity between strains V8 and T16, and also between T14, T15 and T19. Yet, SNPs and indels identified, and subsequently biological activity (see Fig. 1E in particular), supported the notion that all were distinct strains. Taxonomy classification was assigned by utilizing the Type Strain Genome Server (TYGS) [59, 60]. Genomes were uploaded to GeneBank (see above) and associated files can be found in GitHub (see link below).

**Figure 1:**
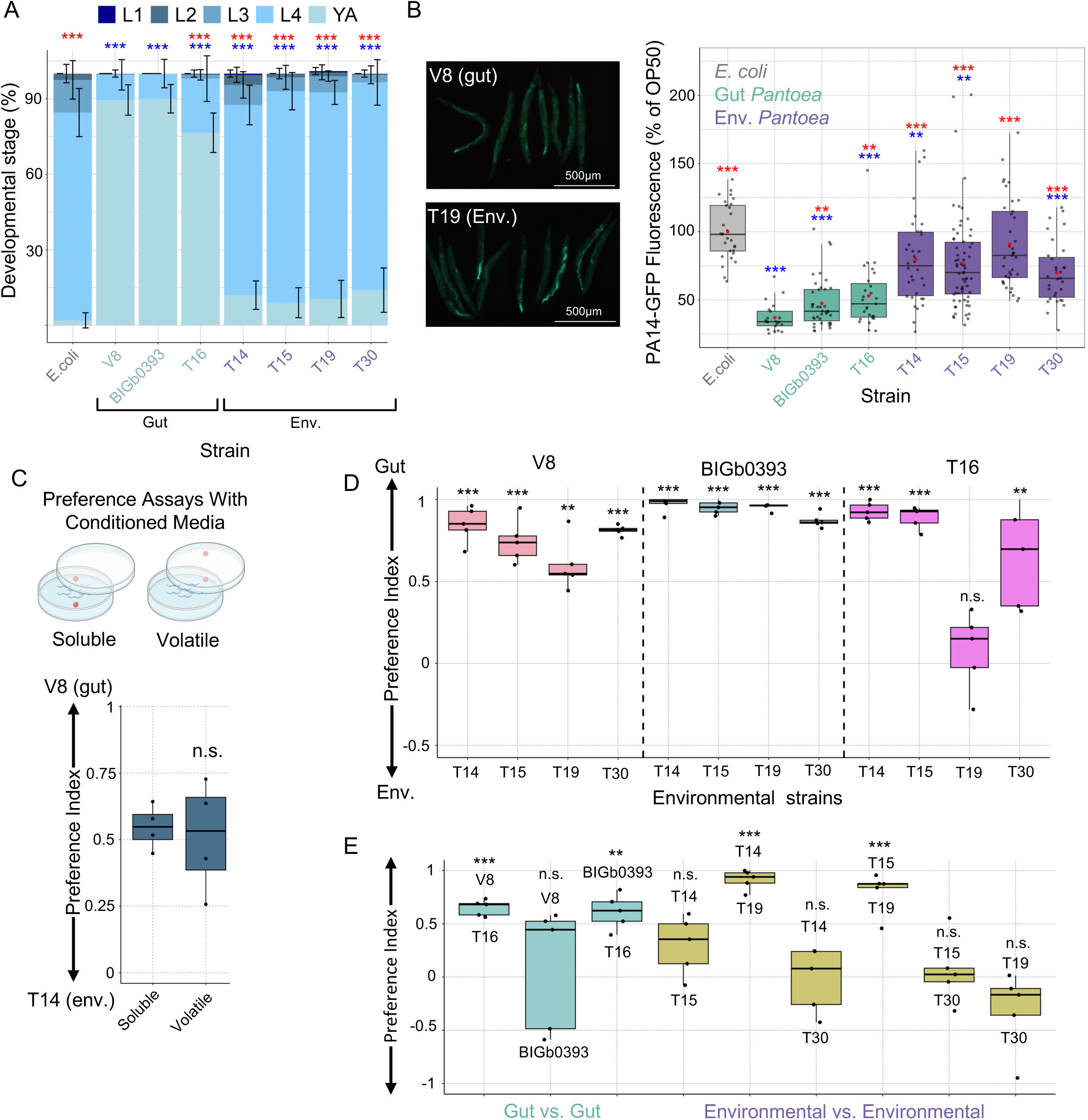
Worms are attracted to the more beneficial *Pantoea* gut isolates. **(A)** Distribution of life stages in populations of developing worms 44 hours after moving synchronized L1 larvae to plates with bacterial strains as designated. Shown are averages ± SDs for 8 replicates (N=25 worms/replicate) for one representative experiment. *, **, ***, *p* < 0.05, *p* < 0.01 and *p* < 0.001, respectively (Fisher Test), red—compared to V8; blue—compared to *E. coli*. **(B)** Representative images of worms infected with PA14-GFP, and quantification of background- and autofluorescence-subtracted signal in all images. Shown are results from three experiments, standardized to their respective *E.coli* controls and put together with boxes stretching between the 25th to 75th percentiles, lines marking medians, and red dots representing mean values; whiskers mark maximum values and dots representing individual worms. *, **, ***, *p* < 0.05, *p* < 0.01 and *p* < 0.001, respectively (t-test). **(C,D,E)** Preference assays between conditioned media (CM) of the designated bacterial strains. **(C)** Spots in scheme represent CM, spotted on plate agar when examining preference based on solutes, or on the plate’s cover, testing preference based on volatiles **(D,E)**. Preference assays in volatile format. Boxes stretch between the 25th to 75th percentiles, lines mark medians; whiskers mark maximum values. Each box represents 4 (C) or 5 (D, E) replicate plates, each with 50-200 worms. *, **, ***, *p* < 0.05, *p* < 0.01 and *p* < 0.001, respectively, t-test (C), or one-sample t-test against 0 with a Benjamini-Hochberg FDR correction (D, E).

### P. cypripedii gene knockouts

Targeted knockout of candidate genes in *P. cypripedii* strain V8 was achieved using the DNA-editing all-in-one RNA-guided CRISPR–Cas transposase (DART) system as previously described [61, 62]. Briefly, target-specific 32-nt spacers were designed to bind in the first half of the target coding sequence adjacent to a 5′-CC Type I-F protospacer adjacent motif. Spacer oligonucleotides (Table S5) were 5′-phosphorylated, annealed, and ligated into BsaI-digested pBFC0619. Ligation products were transformed into *E. coli* BW29427 (a diaminopimelic acid (DAP) auxotroph) via heat shock, and transformants recovered on LB agar supplemented with 20 μg/mL gentamicin and 0.3 mM DAP. Validated donor strains were then conjugated with V8 as described previously [33, 63]. Transposition events were selected on agar plates containing gentamicin but lacking DAP to counter-select against the donor. The genotypes of the 13 mutant strains were confirmed by PCR using gene-specific primers flanking the insertion sites (Table S6, Fig. S1).

### Chemicals

All chemical standards used were purchased at a minimum of 99% purity and were obtained from Millipore Sigma (Sigma-Aldrich, St. Louis, MO, USA).

### Conditioned Media Preparation

Bacterial cultures were pelleted at 3,800 RPM, and supernatant filtered through a 0.22µm filter. This conditioned media (CM) was stored at 4°C (not more than 4 days).

### Behavioral Choice Assay

L1 larvae were added to NGM plates seeded with *E. coli* strain OP50 and allowed to grow for 2 days at 20°C, until the L4 larval stage. Worms were transferred to 1.5 mL tubes and washed three times with M9. Washed worms were pipetted in a minimal volume of M9 onto the center of 60 mm test plates equidistantly (2 cm) from two 3 µL spots, either of bacterial cultures, CM or pure chemoattractant. When spotting bacterial cultures, no antibiotics were included; in the case of V8 mutants, grown with gentamicin, inoculum cultures were pelleted and resuspended in LB without antibiotics, and then grown overnight before spotting on plates. For volatile assays, the spot was placed directly on the underside of the plate’s lid above the marked 2 cm spot from the center. Once worms began to reach any of the two spots (∼15-30 minutes at room temperature), 1µL of 0.5M sodium azide was added onto both spots to paralyze arriving worms and to facilitate counting. After 1 hour, plates were moved to 4°C, until counting (usually within 1-2 days). A preference index (PI) was calculated as (number of worms on strain A) – (number of worms on strain B) divided by (total number of scored worms) [64].

### Phase Extraction from Conditioned Media

Small organic molecules, suspected to include the volatile attraction signal, were extracted from conditioned media using dichloromethane (DCM). CM was mixed with an equal volume of DCM and vigorously shaken. The aqueous layer was aspirated, and residual water in the organic extract removed, first by addition of saturated NaCl brine and then by precipitation with sodium sulfate, filtering-out the precipitate by passage through glass wool. Cleared extract was more strongly preferred by worms compared to the original CM (Fig. S2). Samples of all isolates were processed simultaneously.

### Concentration of DCM Extracts Prepared from Conditioned Media

When in small volume, DCM extracts were concentrated by lightly blowing air on the solution from an air outlet (leveraging the high volatility of DCM compared to any potential solute). When in large volume, a rotary evaporator was used with minimal vacuum (400 mbar, 30°C, ∼1 hour). Following both procedures, a yellow oil formed, which was resuspended in methanol for downstream applications. Concentrated extracts were more strongly preferred by worms compared to the original CM or to the unconcentrated DCM extract (Fig. S2). Samples of all isolates were processed simultaneously.

### HPLC Fractionation and Assessment of Biological activity

Two hundred mL of V8 CM were phase-extracted with 600 mL of DCM and concentrated to ∼20µL on a rotary evaporator. The concentrate was then resuspended in 200 µL MeOH. Fifty µL were then injected onto a C18 column (Phenomenex Luna, 250 × 10 mm, 5 µm particle size) and fractionated using an Agilent 1260 series system equipped with a diode array detector (DAD), monitoring wavelengths of 196 nm, 210 nm, and 254 nm. A solvent system of H_2_O/MeOH (A:B) was used at a flow rate of 2.5 mL/min. A linear gradient was established over 60 min from 95:5 to 5:95, followed by a 25 min isocratic hold at 5:95 for as long as it took for all DAD monitored wavelengths to return to baseline. Seventeen fractions were obtained (collected over consecutive 5 min periods), each of which tested in preference assays against CM from strain T14. Biologically active fractions were subsequently analyzed for chemical composition using GC-MS.

### Gas Chromatography-Coupled Mass Spectrometry (GC-MS)

Samples were analyzed using an Agilent 7890A gas chromatography unit equipped with an Agilent 5975C mass spectrometer. A broad-range Zebron Inferno ZB-5HT column was used for biologically active fractions to identify potential attractants. A higher resolution Restek Rtx-5 column was used for full CM samples to help distinguish between different conformers of candidate attractants. In either case, tert-butyl methyl ether (MTBE) was combined with samples as an internal standard. Both columns used the same run method: 2 min isothermal hold at 35 °C, followed by a ramp to 140°C at a rate of 10°C/min (to slowly volatilize molecules of interest) and a quick ramp from 140°C to 250°C at a rate of 100°C/min, followed by a 2 min isothermal hold at 250°C, to clear the column before the next sample. The range for the mass spectrometer was 20-550 m/z for the bioactive fractions and 20-125 m/z for candidate attractant differentiation. Compound identification was achieved based on the National Institute of Standards and Technology (NIST 8) mass spectral library, and confirmed with pure chemical standards, which were further used for quantification.

Headspace GC-MS was employed to focus on volatiles, and was run on the Agilent 7890A/Agilent 5975C combo and the Restek Rtx-5 column using MS scan parameters of 20-550 m/z. A solid-phase microextraction fiber (SPME) (PDMS/Nitinol core, 24 gauge, 100 μm long, Millipore Sigma) was preconditioned in the instrument’s inlet for 30 minutes at 275°C before insertion into a capped 50 mL Falcon tube with 20mL conditioned media heated to ∼55°C. The fiber was allowed to adsorb volatiles from the headspace above the sample for 20 minutes. It was then inserted and manually injected into the GC-MS inlet (set at 225°C) for 2 min. The temperature gradient used was identical to the liquid injections: 2 min isothermal hold at 35 °C, followed by a ramp to 140°C at a rate of 10°C/min, and then a second ramp from 140°C to 250°C at a rate of 100°C/min, and a final 2 min isothermal hold at 250°C.

### Attractant Concentration Quantification

Concentrations of analytes in biological samples were estimated using a standard curve generated from the respective pure chemicals mixed with water and normalizing to an MTBE internal standard (Fig. S3). Concentrations of butyric acid were estimated based on the standard curve prepared with IAA, which is similar structurally and of almost identical molecular weight.

Headspace concentrations were calculated as described in Supplementary Methods. Briefly, MTBE headspace concentrations were calculated by performing a mole (mass) balance on the sampling apparatus (the closed tube and its headspace) applying Henry’s Law. Relative response factors (RRFs) were calculated from the liquid calibration curves used above in conjunction with MTBE headspace concentration to calculate IAA or 2-PE headspace concentrations at the sampling temperature (55°C). Henry’s Law was again used to transform headspace concentrations of the analytes calculated at 55°C to their respective concentration as would be at the biologically relevant temperature of 20°C.

### Paraformaldehyde-Killed Bacteria

Killing of bacteria was achieved as described elsewhere [65]. Briefly, bacteria were killed by one-hour incubation (with shaking) at 37°C with 0.5% paraformaldehyde (PFA), followed by 5 washes with sterile M9 to remove traces of the fixative. Killing was verified by streaking on LB plates.

### Colony Forming Unit (CFU) Counts

Bacterial density in LB cultures was estimated with CFU counts on LB plates.

CFU counts were also used to evaluate gut colonization. Worms raised on designated bacterial strains or communities from L1 were harvested as young adults, washed from plates with M9/0.025% Triton (M9/T) and surface sterilized as described elsewhere [48]. Following five more washes, live bacteria were extracted from worms, as previously described [33] spread on LB agar and grown at 28°C for 48 hours.

Persistence of bacterial colonization was assessed by transferring colonized adult worms (following 5 washes with M9) to NGM plates with PFA-killed *E.coli* for 24 hours before harvesting for CFU counts.

### Development Rate Measurements

Synchronized L1 larva were plated on NGM plates seeded with designated bacterial strains and raised for two days at 20°C, at which point the percentage of adults was determined. Each plate was divided into quadrants, and the stage of the first 25 worms observed in each quadrant was determined.

### Fluorescence Imaging of Infection

L1 larvae were grown at 20°C to L4 on designated strains. Worms were then washed off plates and transferred to slow killing plates seeded with PA14-GFP and incubated at 25°C [66]. Pathogen lawns were supplemented upon worm transfer with the original bacteria after 24 hours in a 1:1 v/v ratio for replenishment. In plates supplemented with IAA, 50 μL of an 18 mM solution were added after transfer of worms to slow killing plates and added again 24 hours later. Following 44-48 hrs from transfer to PA14 plates, live worms were picked off plates, washed twice with M9/T, paralyzed with 25 mM levamisole, and plated on slides for imaging with a Leica MZ16F stereoscope equipped with a QImaging MicroPublisher 5.0 camera. All images were taken with the same settings.

### Image Analysis

Fluorescent signal in images was quantified using the Fiji plugin in ImageJ v2.10/1.53c [67]. As previously described, background average and autofluorescence gray values for each image were subtracted from their respective Integrated Density values and normalized for worm size to calculate the total fluorescent signal per worm [50].

### Statistical Analyses and Graphics

Statistical analyses were performed in R (v 4.4.1) and described in relevant figure legends. The ggplot2 R package was used to generate most plots [68]. Parts of code written in R for data analysis and graphics and python/command-line interface for genomic assemblies were generated using ChatGPT [69] and subsequently verified and refined. All code, raw data and output files can be found at the GitHub link: https://github.com/jpietropaolo579/-Pietropaolo-et-al.-2026-Data-and-Supporting-Files.

## Results

### Worms are preferentially colonized by fitness-promoting *Pantoea*

Previous work characterizing gut commensals from *C. elegans* worms raised in microbially-diverse environments showed that gut-derived *Pantoea* isolates were more beneficial than congenerics isolated from the worms’ environment, promoting faster development and stronger infection resistance [50]. These fitness benefits were independent of strains’ ability to colonize worms, of which all were capable. Moreover, when given a choice, worms reproducibly preferred the more beneficial gut isolates, suggesting that worms can distinguish between bacterial species based on signals associated with potential benefits, ultimately favoring superior commensals. The previous study compared three gut isolates (*P. nemavictus* strain BIGb0393, and *P. cypripedii* strains V8 and T16) with one environmental isolate (*P. phytostiumlans* T14). To expand this analysis, we tested three additional environmental *Pantoeae* isolates T15, T19, and T30, which genome sequencing identified as distinct *P. phytostimulans* strains (Supp. Tables S2-S4). These additional environmental isolates fully recapitulated the previously observed trends, showing gut colonization when in monocultures (but not as strong as with gut isolates)(Fig. S4), and conferring weaker benefits to worm development and infection resistance, while still outperforming *E. coli* (Fig. 1A and B, Fig. S5).

### Worms are attracted to gut *Pantoea* strains by a secreted volatile signal

Our previous work demonstrated that worms were able to prefer the beneficial *Pantoea*, chemotaxing toward them prior to any contact, which suggested the existence of a bacterial attractant. We found that this attractant was secreted, as worms showed preference not only for spotted bacteria, as previously described, but also to the conditioned media (CM) in which bacteria grew (Fig. 1C). Furthermore, we found that this signal was volatile, as worms preferentially crawled toward CM from a gut isolate when spotted on the plate cover rather than on the agar itself (Fig. 1C). Using this volatile-format attraction assay we found that secreted volatiles were responsible for preference of gut *Pantoea* over environmental congenerics (Fig. 1D). While gut isolates were reproducibly preferred over environmental isolates, some hierarchy was observed, in which among gut isolates V8 and BIGb0393 were preferred over T16 (Fig. 1E) and among the environmental isolates T14 and T15 were preferred over T19, but not over T30. This further supported the notion that T15 and T19, which are very close in their genome sequence, were distinct strains, as were V8 and T16. Altogether, these results demonstrated that worms could sense a secreted volatile signal, which resulted in their preference of the more beneficial *Pantoea* strains.

### Identification of isoamyl alcohol as the bacterial attractant

To isolate and identify the volatile bacterial attractant, we preformed bioactivity-guided fractionation, followed by GC-MS (Fig. 2A). Organic molecules in conditioned media from V8 were extracted with dichloromethane, concentrated on a rotary evaporator, resuspended in methanol, and subjected to HPLC fractionation, giving rise to 17 fractions. Fractions were tested for worm preference over T14 CM, identifying the first six as showing some level of attraction, and thus determined to be bioactive (Fig. 2B). Bioactive fractions were analyzed for their chemical composition employing untargeted GC-MS. Only three molecules were identified: DMSO, which appeared mostly in the first fractions, and is likely a system contaminant, 3,1 methyl butanol (isoamyl alcohol, IAA), and 2-phenylethanol (2-PE) (Fig. 2C). Quantification of the peak areas corresponding to IAA and 2-PE, used as proxies for their relative abundance across fractions, showed strong correlation with host preference (Fig. 2D).

**Figure 2:**
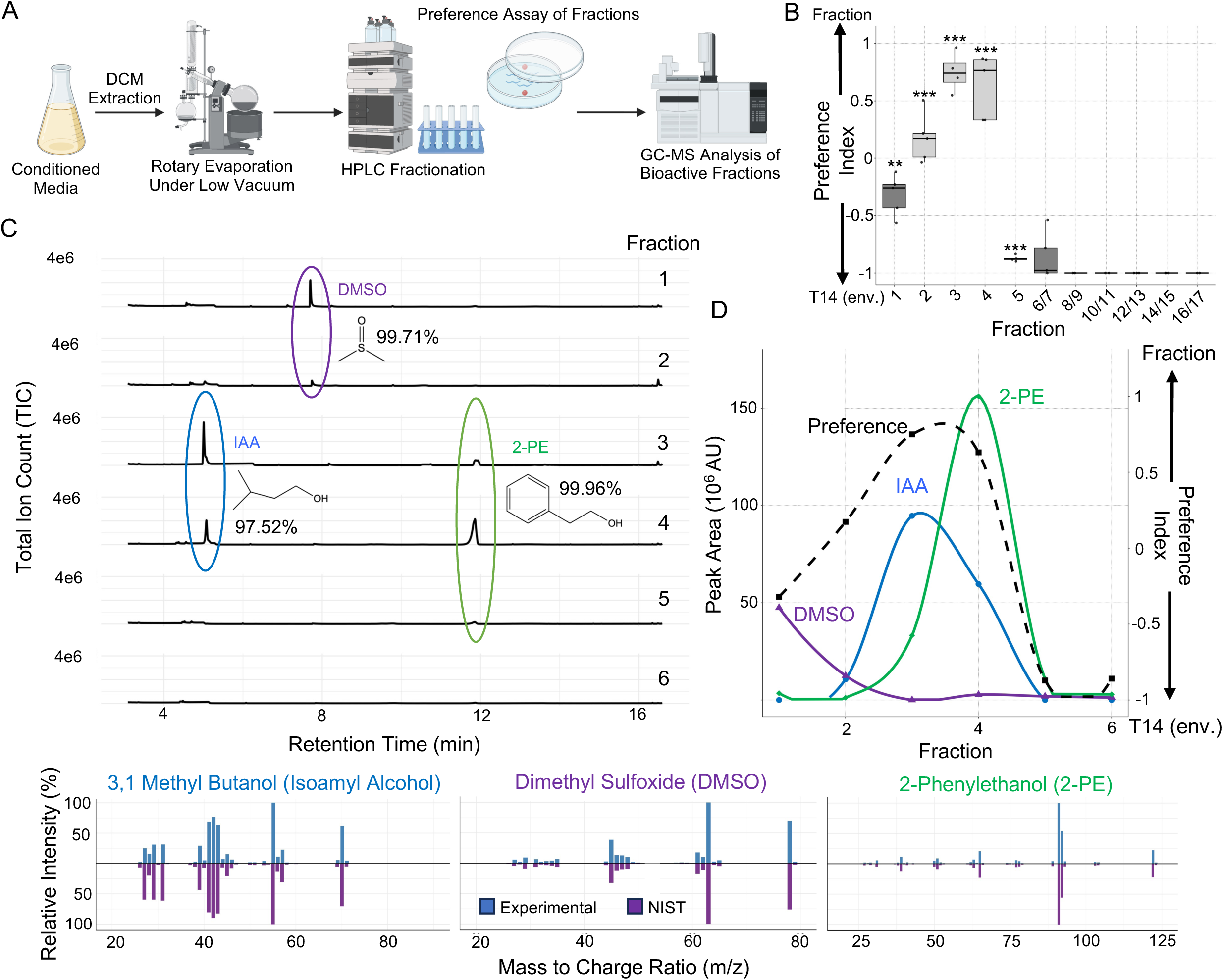
Bioactivity-guided fractionation identifies 3,1 methyl butanol (IAA) and 2-phenylethanol as candidate attractants. **(A)** Purification and characterization pipeline (made with Biorender). **(B)** Worm attraction to HPLC-derived fractions. Boxes stretch between the 25th to 75th percentiles, with lines marking the median; whiskers mark maximum values. Each box represents 5 replicates (plates), each with ∼100-300 worms. *, **, *** indicate *p* < 0.05, *p* < 0.01 and *p* < 0.001, respectively (one-sample t-test compared to -1, which represents solvent only vs. T14). **(C)** GC-MS traces for bioactive fractions with the associated mass spectrum for representative peaks as compared to the NIST database at the bottom. **(D)** Quantification of peak areas for identified metabolites in bioactive fractions, overlaid with average preference index values from (B).

To quantify IAA and 2-PE in conditioned media from different isolates, we performed GC-MS analysis of both conditioned media and the associated headspace (the air directly above the CM in the tube). IAA and 2-PE were detected in all *Pantoea* strains, but not in *E. coli* or in the LB medium (Fig. 3A and B). Concentrations of the two metabolites in CM from the different strains were calculated based on calibration curves generated with pure standards (Fig. S3 and Supplementary Methods) and normalized to the optical density in the original cultures. This analysis overall indicated higher secretion by gut isolates compared to environmental isolates (Fig. 3C and D, top). but differences were not clear-cut. However, normalization of concentrations to the number of viable bacteria, determined by CFU counts (Fig. S6), revealed markedly higher production of both IAA and 2-PE by gut commensals compared to environmental strains (Fig. 3C and D bottom).

**Figure 3.**
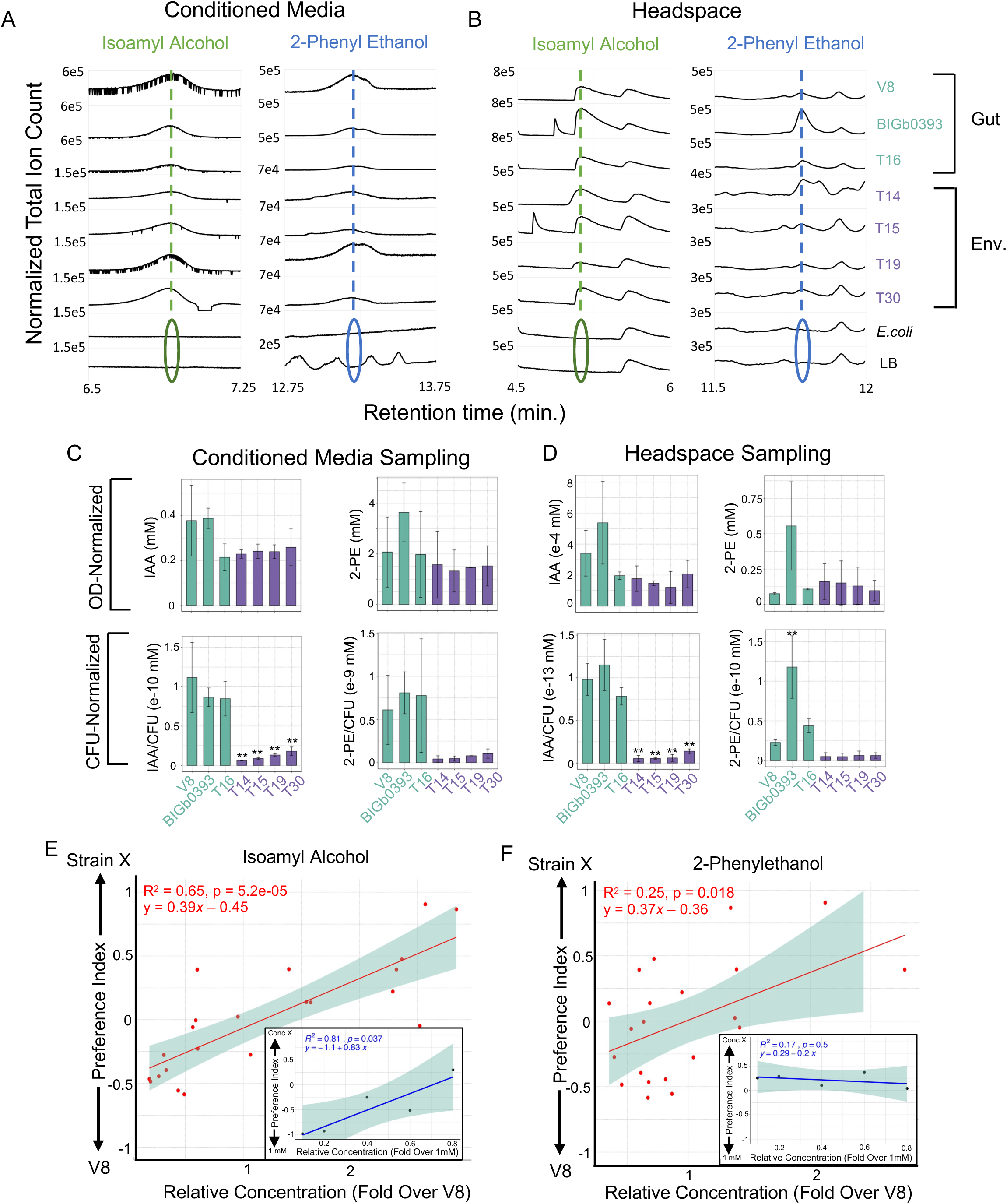
Gut-colonizing *Pantoea* produce much higher concentrations of isoamyl alcohol (IAA) and 2-phenylethanol (2-PE) than non-gut colonizing strains. **(A,B)** GCMS analysis of conditioned media, normalized to an internal standard MTBE, and zoomed in to highlight the relevant metabolites **(A)** or of the media’s headspace **(B)** from LB cultures of designated *Pantoea* strains, with peaks of IAA and 2-PE marked. Note the different y-axis range for samples from gut and environmental isolates. Shown is one representative measurement. **(C,D)** Quantification of IAA and 2-PE concentrations in conditioned media **(C)** and the media’s headspace **(D)** using an internal standard. Shown are averages ± SDs for measurements in 2 independent experiments normalized for culture OD (top) or CFU (bottom). Statistical significance was assessed by one-way ANOVA with Dunnett’s post-hoc test comparing each strain to V8; *, **, ***, *p* < 0.05, *p* < 0.01 and *p* < 0.001, respectively. **(E,F)** Correlation between concentrations of IAA (E), 2-PE (F) in CM and worm preference of the same CM. Concentrations were calculated using a standard curve and normalized to an internal standard. Shown are best fit lines and standard errors (shade) and the respective line formulas, correlation coefficient and statistical significance (Pearson correlation). **Insets** describe similar correlations with the respective metabolites in pure form.

To test whether worm preference of the different isolates correlated with levels of secreted IAA or 2-PE, we compiled preference indices from multiple independent volatile-format assays comparing CM form different strains to V8 CM, together with metabolite concentrations measured in the corresponding CMs samples by GC-MS as in Fig. 3C (OD-normalized). This analysis revealed a significant correlation between worm preference and secreted levels of either IAA or 2-PE (Figs 3E and F). In contrast, similar analyses using defined concentrations of pure IAA or 2-PE within the range detected in bacterial CM (0.1-0.5 mM for IAA, 0.5-5 mM for 2-PE) showed a significant correlation with worm preference only for IAA, whereas 2-PE concentrations showed no significant correlation (Fig. 3E,F insets). Together, these experiments support a central role for secreted IAA levels in determining *C. elegans* preference of *Pantoea* strains, while suggesting that 2-PE is unlikely to function as an attractant itself, at least within the tested concentration range (see Fig. S7), and may simply correlate with IAA secretion.

### Sensing of IAA from beneficial *Pantoea* guides preferred gut colonization

The identification of IAA as the metabolite mediating worm attraction to specific *Pantoea* strains connected our findings to a large body of work in which IAA has served as a model compound for studying neuronal control of *C. elegans* behavior. To gain better understanding of host genes involved in sensing IAA at concentrations relevant for commensal preference we examined preference of IAA by mutants previously implicated in chemotactic response to IAA, using IAA concentrations comparable to those secreted by *Pantoea* isolates. This analysis identified *odr-1, 2* and *5*, and *sra-11* mutants as significantly deficient in sensing the attractant (Fig. 4A).

**Figure 4:**
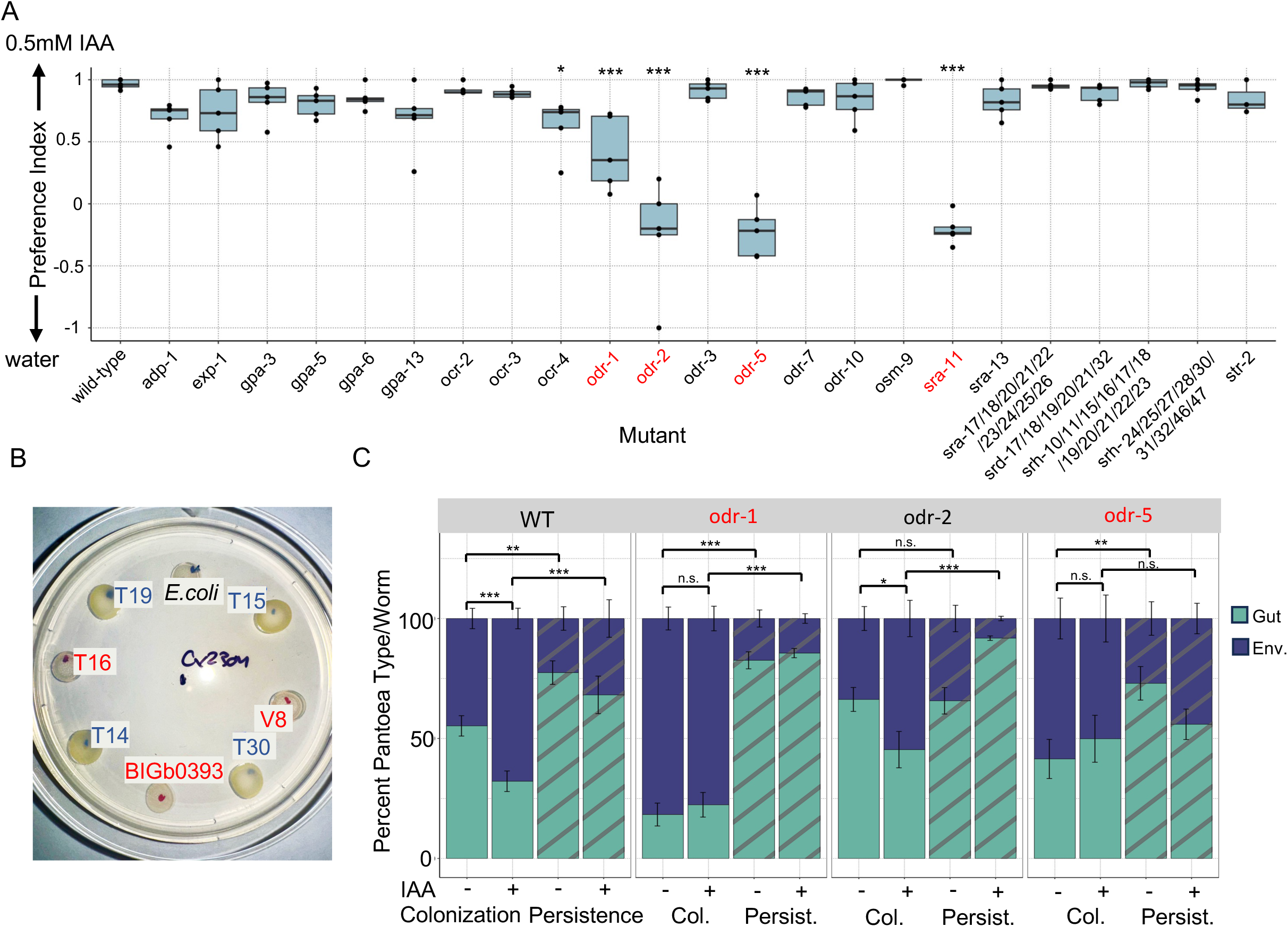
IAA sensing is important for determining gut colonization. **(A)** Volatile-format preference assays for strains disrupted for IAA-sensing candidate genes. Boxes stretch between the 25th to 75th percentiles, with lines mark medians; whiskers mark maximum values. Each box represents 5 technical replicates (plates), each with ∼100-300 worms. *, **, *** indicate *p* < 0.05, *p* < 0.01 and *p* < 0.001, respectively, one-way ANOVA with Dunnett post-hoc. **(B)** A representative spatially segregated culture plate on which L1s were raised until the gravid stage at which point they were washed, ground, and their gut microbiome composition analyzed by CFU counts. Environmental *Pantoea* cultures in blue were treated with IAA (+) or left untreated (-); Gut *Pantoea* strains, in red, and *E.coli* were not supplemented with IAA. **(C)** Colonization bars show the percentage of CFUs of gut *Pantoea* of the two types, in gravid worms raised on the spatially segregated plates (as in B). Shown are averages ± standard errors for n=18-30 counts/group (2 independent experiments, each with 9-15 technical replicates), representing a total of 66-199 worms/group (median = 116 worms). Bars marked as Persistence represent microbiome composition in similarly raised worms shifted after colonization to plates with only PFA-killed *E. coli* for additional 24 hours. *, **, *** indicate *p* < 0.05, *p* < 0.01 and *p* < 0.001, respectively, Wilcox ranked-sum t-test.

*odr-1* encodes a guanylate cyclase expressed in multiple neurons, including the amphid neurons AWB and AWC, which, together with AWA, are the central regulators of attraction to volatile chemicals [70] *odr-2* encodes a Ly-6-related glycosylphosphatidylinositol (GPI)-anchored extracellular membranal protein that is expressed in many sensory neurons, including AWC amphids [71]*. odr-5* encodes a protein of unknown function expressed primarily in AWC neurons [72], whereas *sra-11* encodes a G-protein-coupled receptor (GPCR) previously studied in interneurons associated with AWA- and AWC-mediated responses [73].

Using the identified mutants, we tested the importance of IAA sensing for gut colonization. The seven *Pantoea* isolates were spotted on several plates in random order, together with the non-colonizing *E. coli*. L1 larvae were placed at the center of the plates and allowed to forage as they developed to adulthood (Fig. 4B). In parallel experiments, IAA was added to spots containing environmental strains in order to uncouple strain identity from IAA secretion. After three days, adult worms were harvested, surface-sterilized, washed, and homogenized, and gut bacteria were quantified by CFU counts, distinguishing gut-derived and environmental isolates based on the yellow-orange appearance of the latter (Fig. 4B). Wildtype animals showed preferential colonization by gut-derived strains (Fig. 4C, Fig. S8). However, supplementation of environmental strains with IAA abolished this preference, leading to increased representation of environmental strains in the gut. Two of three tested neuronal mutants - *odr-1* and *odr-5* - were defective in preferential colonization by high IAA-secreting gut strains even in the absence of exogenous IAA, and addition of IAA to environmental strains did not further alter their colonization patterns. In contrast, *odr-2* mutants displayed colonization patterns similar to those of wildtype animals. This may reflect a broader defect in sensing bacterial or food-associated cues rather than a specific defect in IAA sensing, as *odr-2* mutants in preference assays (Fig. 4A) failed to accumulate near IAA spots altogether and instead were scattered evenly throughout the plate (Fig. S9A). In addition, these mutants were substantially slower than wildtype worms or other mutants in locating food sources (Fig. S9B). Together, these experiments support a key role for IAA sensing in the preferential uptake and colonization by specific commensal strains.

Although IAA sensing was important for initial acquisition of commensals, further investigation revealed an additional layer of selection within the gut. When worms were shifted after initial colonization to plates containing only food *E. coli* (PFA-killed) and maintained for an additional day in the absence of *Pantoea* bacteria, gut-derived isolates persisted and expanded within the intestine, ultimately dominating over environmental strains regardless of their initial proportions at the time of transfer. This suggested that in addition to host sensing and preference of beneficial strains, gut-derived commensals possessed intrinsic competitive advantages that enabled them to outcompete other strains and establish dominance within the gut niche independently of host behavior.

### An *ilvE* Homolog Contributes to IAA Production in *Pantoea*

To investigate IAA synthesis in *Pantoea*, we sought to identify bacterial mutants impaired in IAA production. In bacteria, IAA can be produced from leucine through the bacterial equivalent of the Ehrlich pathway [74, 75]. Examination of the V8 genome identified thirteen genes encoding enzymes with potential roles in this pathway, including aminotransferases, α-keto acid decarboxylases and alcohol/aldehyde dehydrogenases (Fig. 5A, Table S7). Knockout mutants were generated for each candidate gene and tested in worm preference assays against wildtype V8. This screen identified *ilvE1*, a homolog of a branched-chain amino acid aminotransferase, as a candidate contributor, as its disruption reduced worm preference relative to wildtype V8, although the effect did not reach statistical significance (Fig. 5B). Supporting a role for *ilvE1* in IAA production, CM from the Δ*ilvE1* mutant - the same samples used in the preference assays - showed significantly reduced IAA concentrations (Fig. 5C). Interestingly, GC-MS traces of CMs from the Δ*ilvE1* mutant identified a metabolite peak that increased dramatically compared to wildtype V8 (Fig. 5D). Analysis of its mass spectrum identified it as butyric acid (BA), the protonated form of butyrate, and its quantification showed a 5-fold increase in its concentration in Δ*ilvE1* CM compared to wildtype, without any large change in CMs of other mutants. Which *Pantoea* pathway is responsible for producing BA is unknown, but its increase may represent a shift in the metabolic or energetic flux from IAA to BA associated with the Δ*ilvE1* disruption. Altogether, the results suggest that *ilvE1* contributes to IAA biosynthesis in *Pantoea*. However, the modest effect of its disruption on worm preference, as well as the continued secretion of detectable IAA by the mutant, suggests that IAA synthesis in *Pantoea* may involve substantial pathway redundancies.

**Figure 5:**
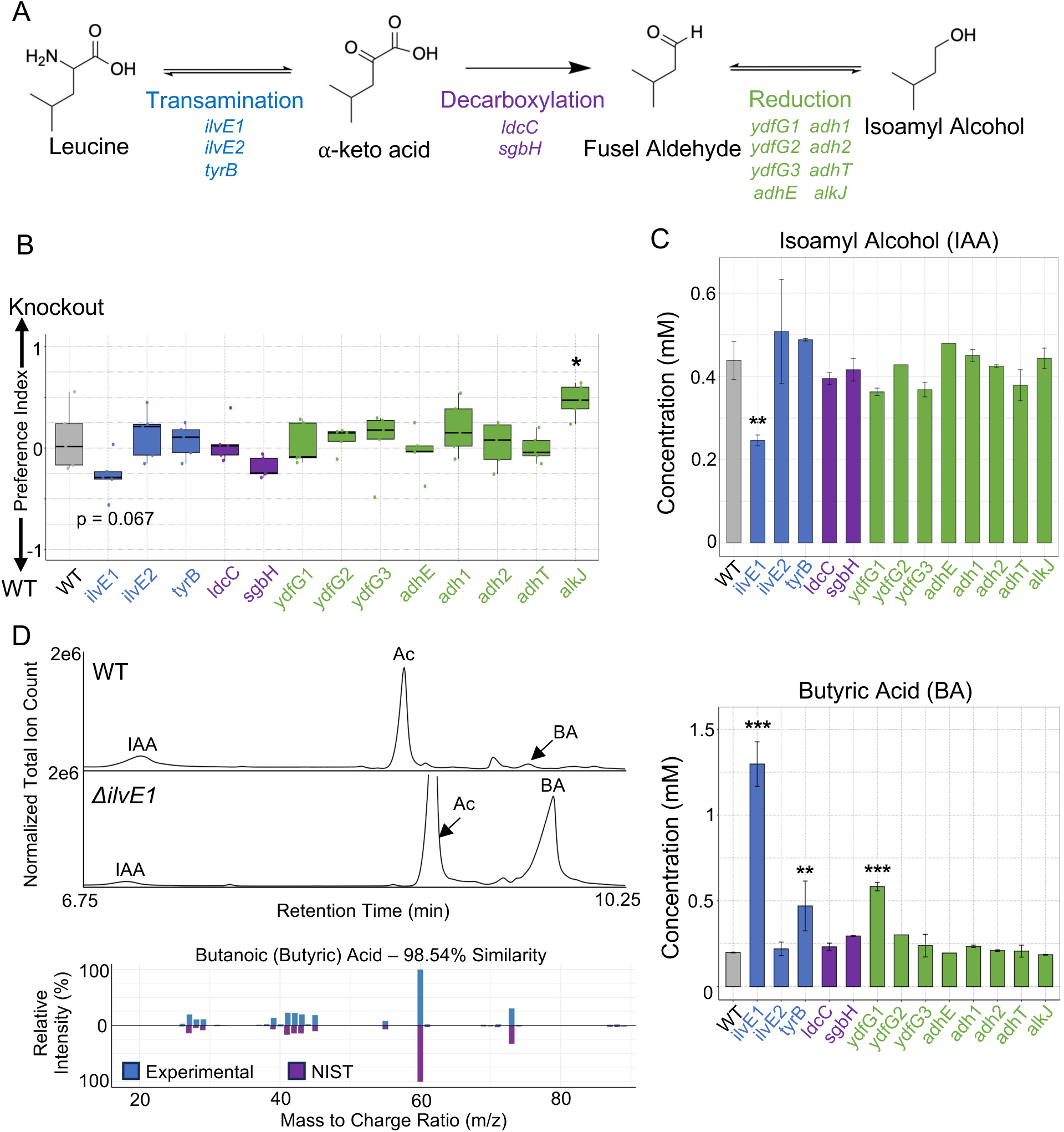
Knockout of the V8 strain branched chain amino acid transferase *ilvE1* results in decreased preference for supernatant and lower isoamyl alcohol production. **(A)** The proposed pathway leading from leucine to IAA production, and candidate V8 enzymes that may take part in it (identified based on PROKKA annotations). **(B)** Worm preference of V8 mutants vs. wildtype. Boxes stretch between the 25th to 75th percentiles, with lines mark medians; whiskers mark maximum values. Each box represents 5 technical replicates (plates), each with ∼100-300 worms. (**C**) IAA concentrations in mutants. **(D)** Left, representative GC-MS traces for WT and *ΔilvE1* CM showing peaks for IAA and for butyric acid (BA, the protonated form of butyrate), with the associated mass spectrum for a representative peak as compared to the NIST database; also shown in traces is a peak identified as acetic acid (acetate), which did not show significant differences among samples. Right, quantified concentrations of BA, averages ± SDs from two measurements. One-way ANOVA followed by Dunnett’s post-hoc was employed in both C and D; *, **, ***, *p* < 0.05, *p* < 0.01 and *p* < 0.001, respectively.

**Fig. 6.**
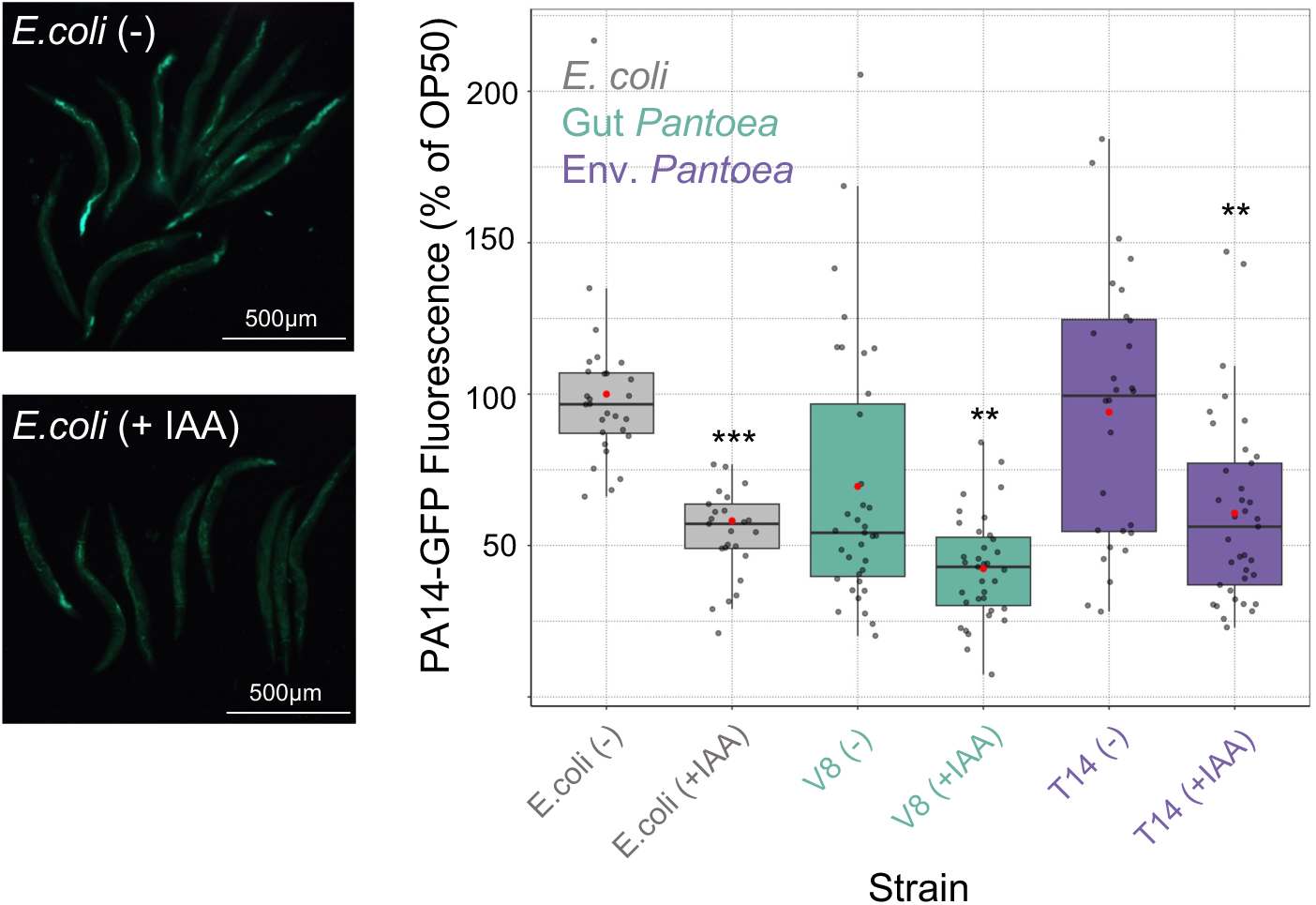
Isoamyl alcohol promotes pathogenic infection resistance. Representative images of worms infected with PA14-GFP, and quantification of background- and autofluorescence-subtracted signal in all images. Shown are results from three experiments, standardized to their respective *E.coli* controls and put together with boxes stretching between the 25th to 75th percentiles, lines marking medians, and red dots representing mean values; whiskers mark maximum values and dots representing individual worms. *, **, ***, *p* < 0.05, *p* < 0.01 and *p* < 0.001, respectively (t-test).

### IAA Contributes to Host Fitness

Recent work has proposed that IAA may serve an indicator of nutrient-rich bacteria or environments [47]. We asked whether IAA may contribute also directly to host fitness. Focusing on the ability of *Pantoea* V8 to limit pathogenic colonization, we found that worms raised on bacterial lawns supplemented with IAA exhibited further reduction in pathogenic colonization by PA14-GFP compared to worms raised on unsupplemented lawns (Fig. 5). This effect was observed in worms raised on V8, T14 and on non-colonizing, non-IAA producing *E. coli*, suggesting that IAA itself can enhance host protection against pathogen colonization and thereby improve fitness. Together, these findings indicate that IAA may provide a direct benefit to the host, beyond serving merely as a proxy for nutrient availability.

## Discussion

Our findings identify isoamyl alcohol (IAA) as a secreted volatile bacterial signal that links microbial amino acid metabolism to host acquisition of beneficial commensals from a microbially complex environment. We show that *C. elegans* senses IAA, likely via sensory amphid neurons, and chemotax preferentially toward *Pantoea* strains with higher IAA secretion. Notably, these high-secreting strains are also more beneficial to the host. This behavioral preference facilitates early colonization with superior commensals, ultimately increasing host fitness. These results suggest that host behavior plays a role in shaping gut microbiome composition by promoting selective acquisition of specific bacteria. Furthermore, the host ability to detect subtle, strain-level metabolic differences carries significant eco-evolutionary consequences. While IAA may serve as a proxy for the essential amino acid leucine, we found that on its own, added to different bacterial species, IAA enhanced host infection resistance and fitness, suggesting that it modified not only host behavior, but also its physiology. More broadly, our results imply that animals may exploit bacterial metabolic byproducts as proxies for nutrient availability, microbial physiology, and symbiotic potential.

IAA has long served as a model attractant in *C. elegans* neuroscience research, frequently used to interrogate the mechanisms of neuronal sensing and behavioral regulation [76, 77]. More recently, IAA gained attention as a relevant bacterial metabolite [78] and a putative signal for essential amino acids, being a product of leucine catabolism [47]. Our results extend these ideas by demonstrating that IAA is also a signal for beneficial commensals. Furthermore, the ability of IAA to independently increase host fitness suggests a role in modulating host physiology. While what this role may be is yet unknown, a previous study reported that IAA-exposed *C. elegans* exhibited increased stress tolerance and longevity, supporting our findings [79]. These findings also align with earlier results showing that dead BIGb0393 *Pantoea* protected worms as effectively as live ones, in agreement with a metabolite-mediated protective effect [50].

Leveraging the extensive knowledge on IAA sensing, we identified several genes that were important for distinguishing *Pantoea* strains. Among our candidates, *odr-1* and *odr-5* were the most prominent. Both are expressed primarily in amphid sensory neurons, specifically in AWC neurons [72, 80]. Mutants impaired in IAA sensing lost their preference for gut-associated *Pantoea* or supplemented IAA. Conversely, supplementing low-secreting *Pantoea* with IAA was sufficient to “mislead” wildtype worms into preferentially acquiring inferior commensals. These findings establish IAA sensing as a dominant factor in determining choice of commensals and in microbiome assembly.

While behavioral preference for higher IAA concentrations was dominant in determining initial gut colonization, *Pantoea* strains naturally high in IAA were able to outcompete environmental strains in the gut even when initially outnumbered and without further uptake (Fig. 4C). Disruption of IAA sensing altered early colonization patterns but did not prevent gut-associated strains from ultimately dominating the gut niche. This suggests that long-term colonization is determined by two distinct factors: initial recruitment driven by host sensing and behavior, and establishment of commensalism, driven by bacterial competitive fitness within the intestinal environment. This duality mirrors recent work in *C. elegans* showing that microbiome remodeling that enabled host resistance to an environmental toxin was driven independently by host preference toward toxin-modifying bacteria and by the increased competitive advantage of the protective bacteria [65]. Together, these studies support a multi-step model of commensal acquisition and microbiome assembly involving both host factors and microbial adaptations.

We identified *ilvE1* as important for IAA biosynthesis. While its disruption weakened worm attraction, it did not abolish IAA secretion entirely, suggesting biochemical redundancy. Supporting this, a previous report demonstrated that *tyrB*, normally an aromatic amino acid aminotransferase, could promiscuously modify, in *ilvE*-disrupted *E.coli,* branched chain amino acids [81]. Similar redundancies may act elsewhere in the pathway. How this translates to differences in the secreted levels of IAA between gut commensals and environmental *Pantoea* strains remains unknown, but could involve sequence divergence among the different enzymes, and/or differences in their expression.

The finding that worms are not attracted to 2-PE at the concentrations secreted by *Pantoea* was somewhat surprising, given that 2-PE, similar to IAA, is derived from the essential amino acid (EAA) phenylalanine. Thus, one might expect it to be a significant nutritional signal, as IAA. However, there may be a hierarchy of importance among EAAs; for instance, only five EAAs out of ten - including leucine, but not phenylalanine - were previously shown to extend *C. elegans* lifespan [82]. IAA may therefore be a higher-priority signal. Furthermore, the independent contribution of IAA to host fitness may increase selective pressure to maintain sensitive IAA detection. Thus, while 2-PE levels correlate with IAA levels - possibly due to shared metabolic enzymes or pathway regulators - 2-PE alone is insufficient to drive host preference.

In summary, our findings support a model in which *C. elegans* leverages metabolite sensing to selectively acquire beneficial microbes, with subsequent gut persistence determined by bacterial competitive traits. This study highlights behavioral selection based on sensory cues as an important yet underexplored factor in microbiome assembly and host fitness. Our findings suggest a general mechanism where active host behavior shapes microbial landscapes, complementing immune and nutritional filtering.

## Acknowledgements

We thank McKenna Yao and Wenjun Zhang of UC Berkeley’s Chemical and Biomolecular Engineering department for allowing us to use their HPLC and for their advice on headspace chemical concentration calculations. We also thank Yi Liu from the EBI facility at UC Berkeley for GC-MS troubleshooting.

## Conflicts of Interest

The authors declare no conflicts of interest.

## Data Availability

All code and data files can be found at the GitHub link: https://github.com/jpietropaolo579/-Pietropaolo-et-al.-2026-Data-and-Supporting-Files.

## Funding

Work described in this manuscript was supported by NIH grants (received by MS) R01OD024780 and R01AG061302.

---

**Figure S1:**
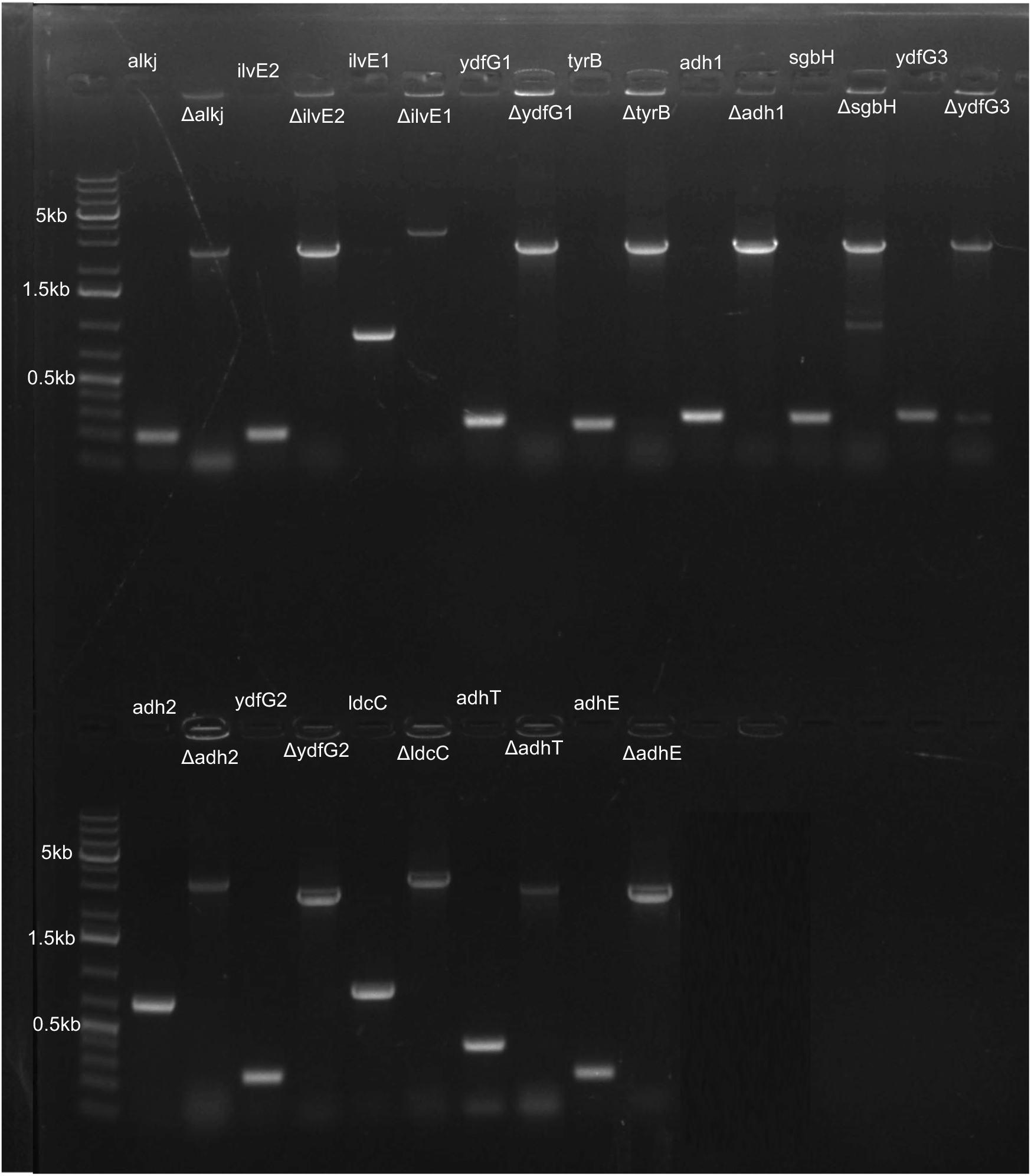
Verification of V8 knock-out mutants. Shown are amplification products for wildtype and knock-out alleles of targeted genes.

**Figure S2.**
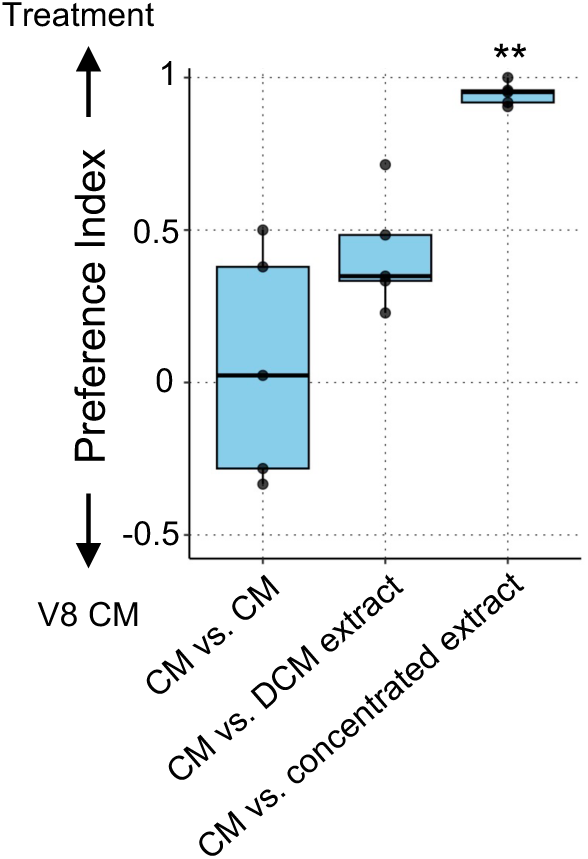
Comparison of worm attraction to *Pantoea* conditioned media and its extracts. Boxes stretch between the 25th to 75th percentiles, with lines mark medians; whiskers mark maximum values. Each box represents 5 technical replicates (plates), each with ∼100-300 worms. *, **, *** indicate *p* < 0.05, *p* < 0.01 and *p* < 0.001, respectively (t-test).

**Figure S3:**
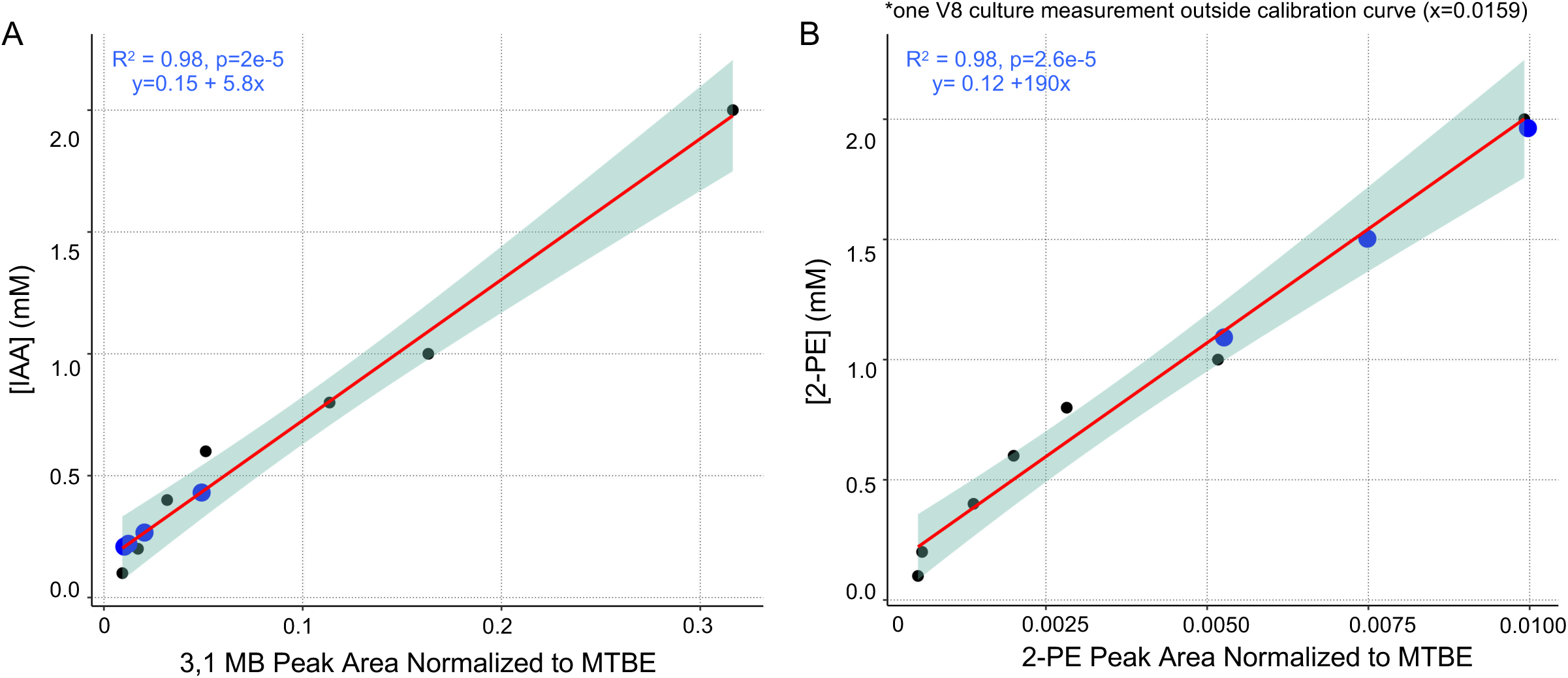
Standard curves for estimating concentrations from MTBE-normalized peak areas. Curves for isoamyl alcohol (IAA) (**A**) and 2-phenylethanol respectively (2-PE) (**B**). Black dots represent the pure chemical in known concentrations used to construct the curves. Blue dots represent measurements of MTBE-normalized peak areas of the respective molecule in V8 culture samples (measured across all experiments). Shown also are best-fit lines (red), standard errors (green shading), and statistical significance (Pearson correlation).

**Figure S4:**
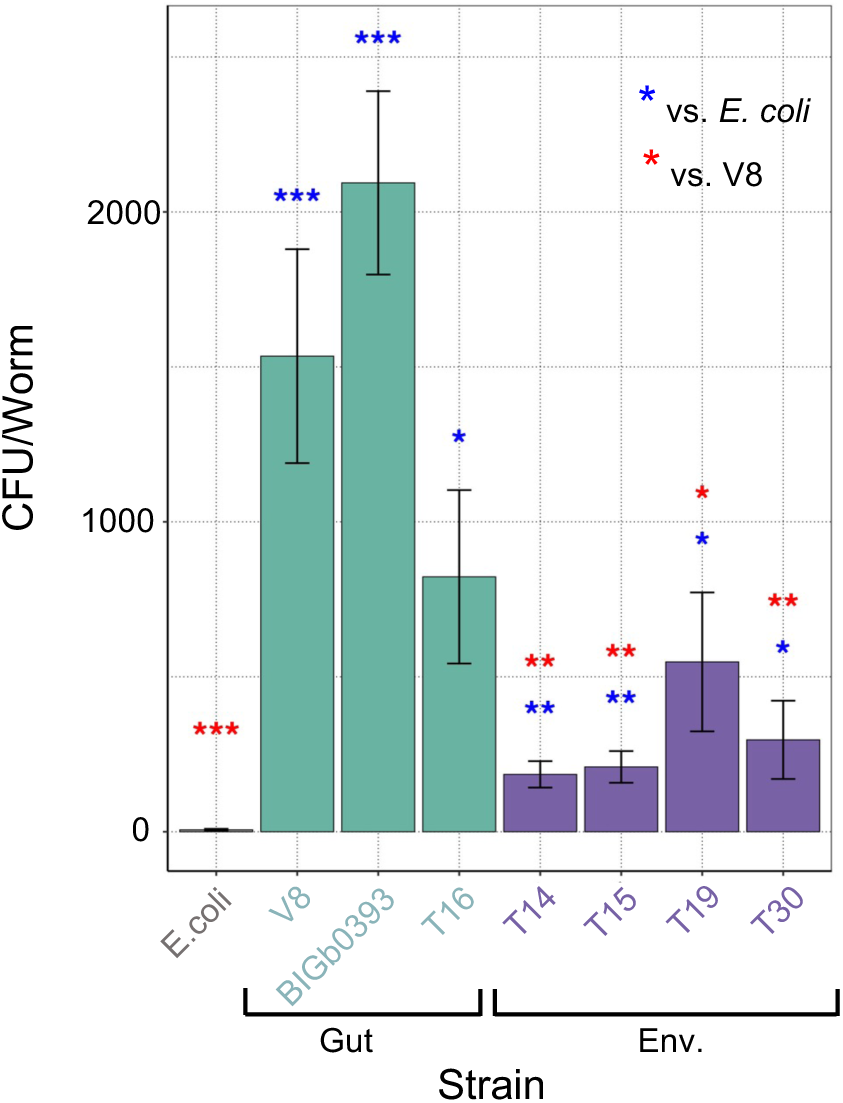
All examined *Pantoea* strains can colonize worms when in monocultures. Worm gut CFU counts in adult worms raised on monocultures of designated strains for three days. Bars averages ± standard deviations for n=18 counts per group (2 independent experiments, each with 9 technical replicates), representing a total of ∼72-190 worms/group (median ∼ 146 worms). *, **, *** indicate *p* < 0.05, *p* < 0.01 and *p* < 0.001, respectively (t-test).

**Figure S5:**
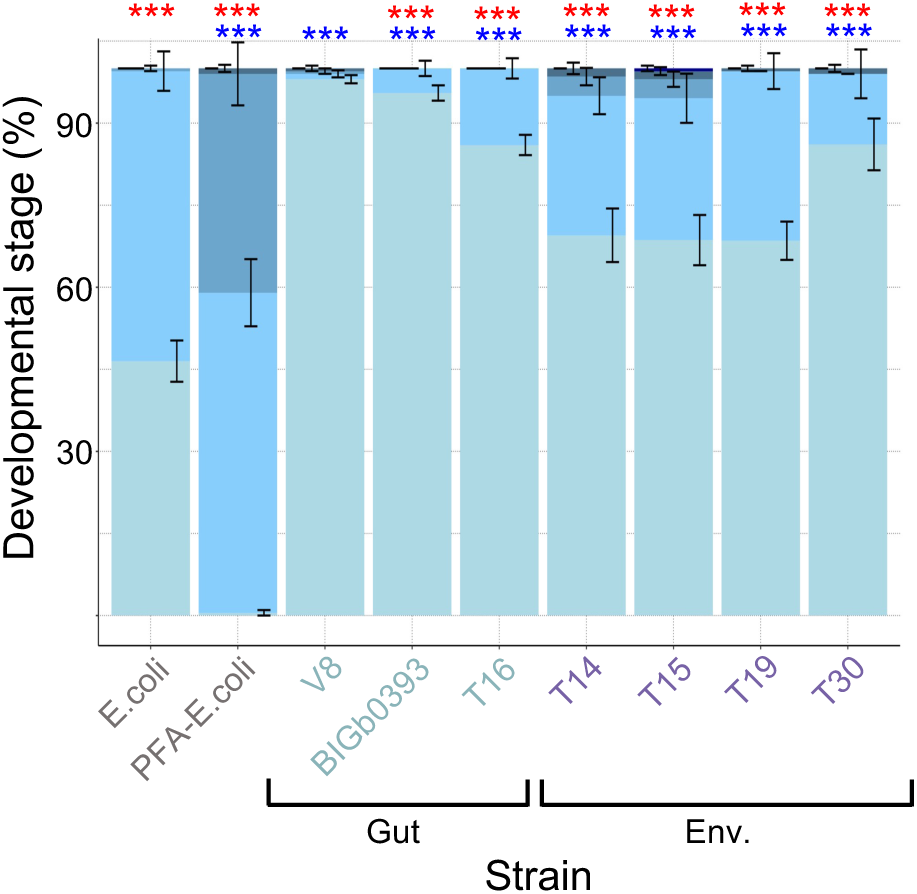
An independent repeat of the experiments presented in Fig. 1A demonstrating faster worm development with *Pantoea* gut isolates than with environmental *Pantoea* strains. Distribution of life stages in populations of developing worms 44 hours after moving synchronized L1 larvae to plates with designated bacterial strains. Shown are averages ± SDs for 8 replicates (N=25 worms/replicate). *, **, ***, *p* < 0.05, *p* < 0.01 and *p* < 0.001, respectively (Fisher Test), red - compared to V8; blue - compared to *E. coli*.

**Figure S6:**
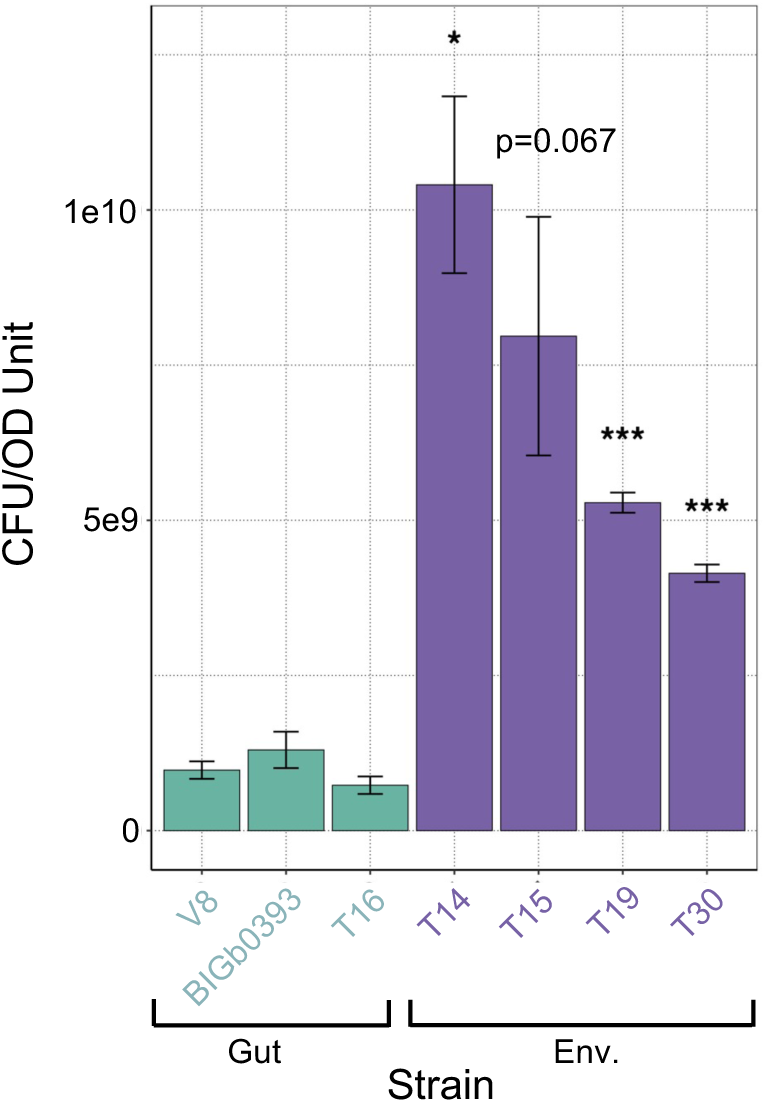
CFU/OD ratio is higher for cultures of environmental isolates compared to gut isolates. Bars represent CFUs from 3 independent LB cultures, shown as averages ± standard deviation. *, **, *** indicate *p* < 0.05, *p* < 0.01 and *p* < 0.001, respectively (t-test compared to V8).

**Figure S7:**
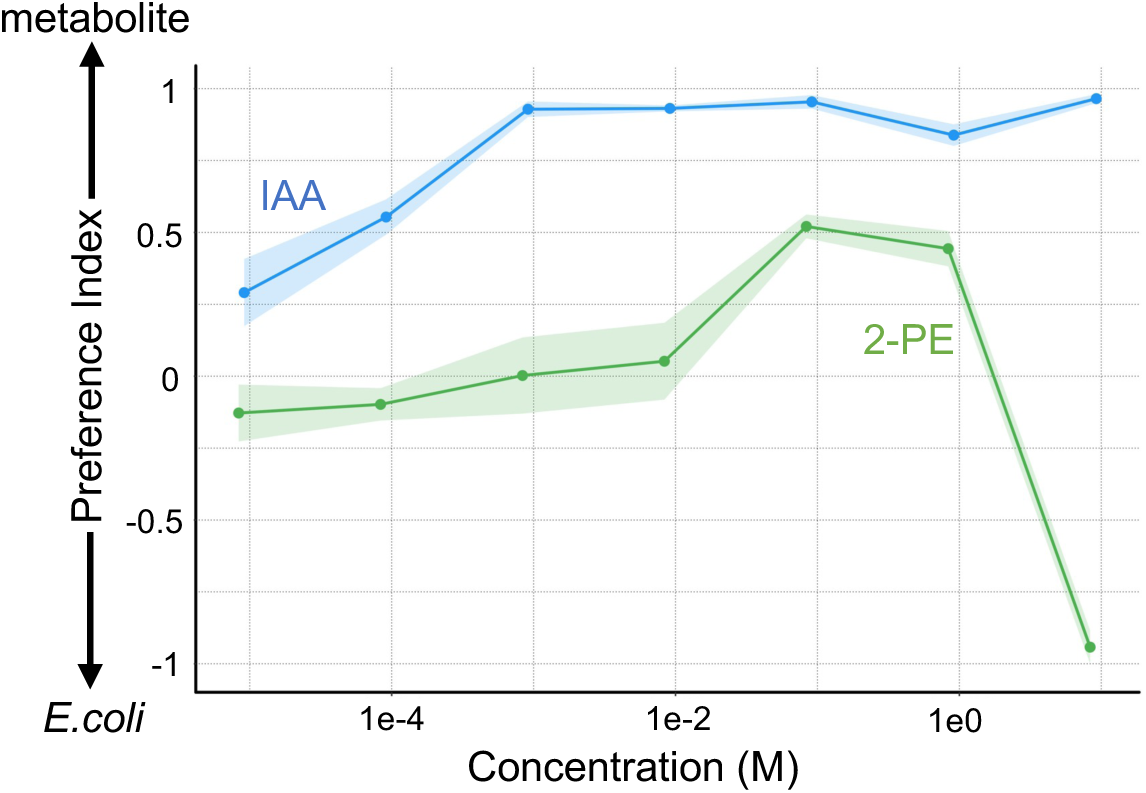
IAA and 2-PE are preferred by worm over E. coli in different concentration ranges. A preference index of 1 represents a preference for the respective metabolite. Points represent an average of counts from 5 plates (n=50-100 worms per plate) used in a volatile-format preference assay between pure metabolites in the designated concentration vs. *E.coli* conditioned media. Shading represents standard error. Note the higher concentrations required for preference of 2-PE (10-100 mM) compared to the lower concentrations in which IAA functions (10mM-1mM)

**Figure S8:**
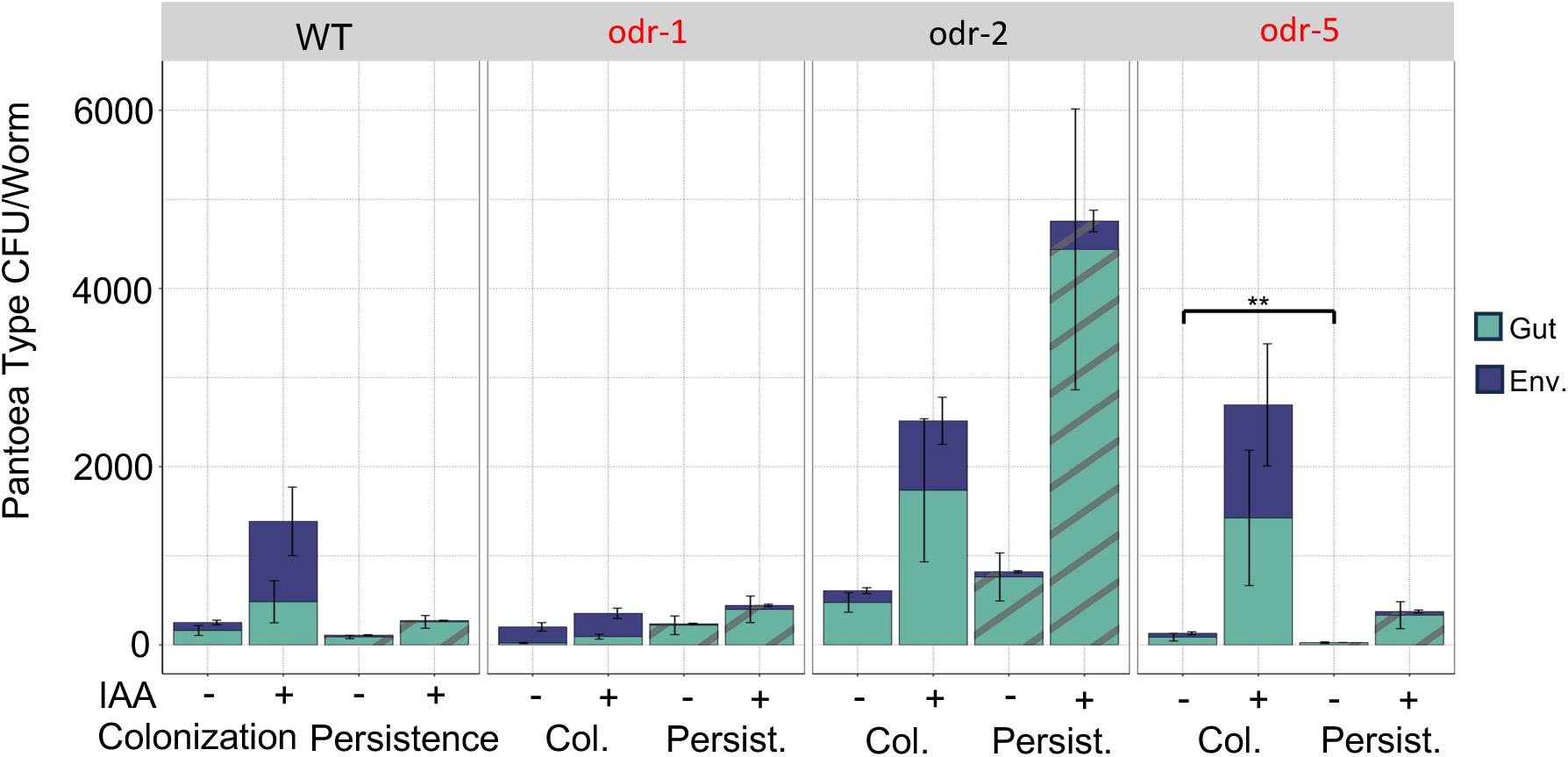
Total CFU counts of gut *Pantoea* shown as percentages in. **Figure 4C**. Bars represent CFUs of gut *Pantoea* of the two designated types. Shown are averages ± standard errors for n=18-30 counts per group (2 independent experiments, each with 9-15 technical replicates), representing a total of 66-199 worms/group (median = 116 worms). Colonization bars represent microbiome composition in gravid worms raised on the spatially segregated plates (as in Fig. 4B). Persistence bars represent microbiome composition in similarly raised worms shifted after colonization to plates with only PFA-killed *E. coli* for additional 24 hours. ** indicate *p* < 0.01, Wilcox ranked-sum t-test.

**Fig. S9.**
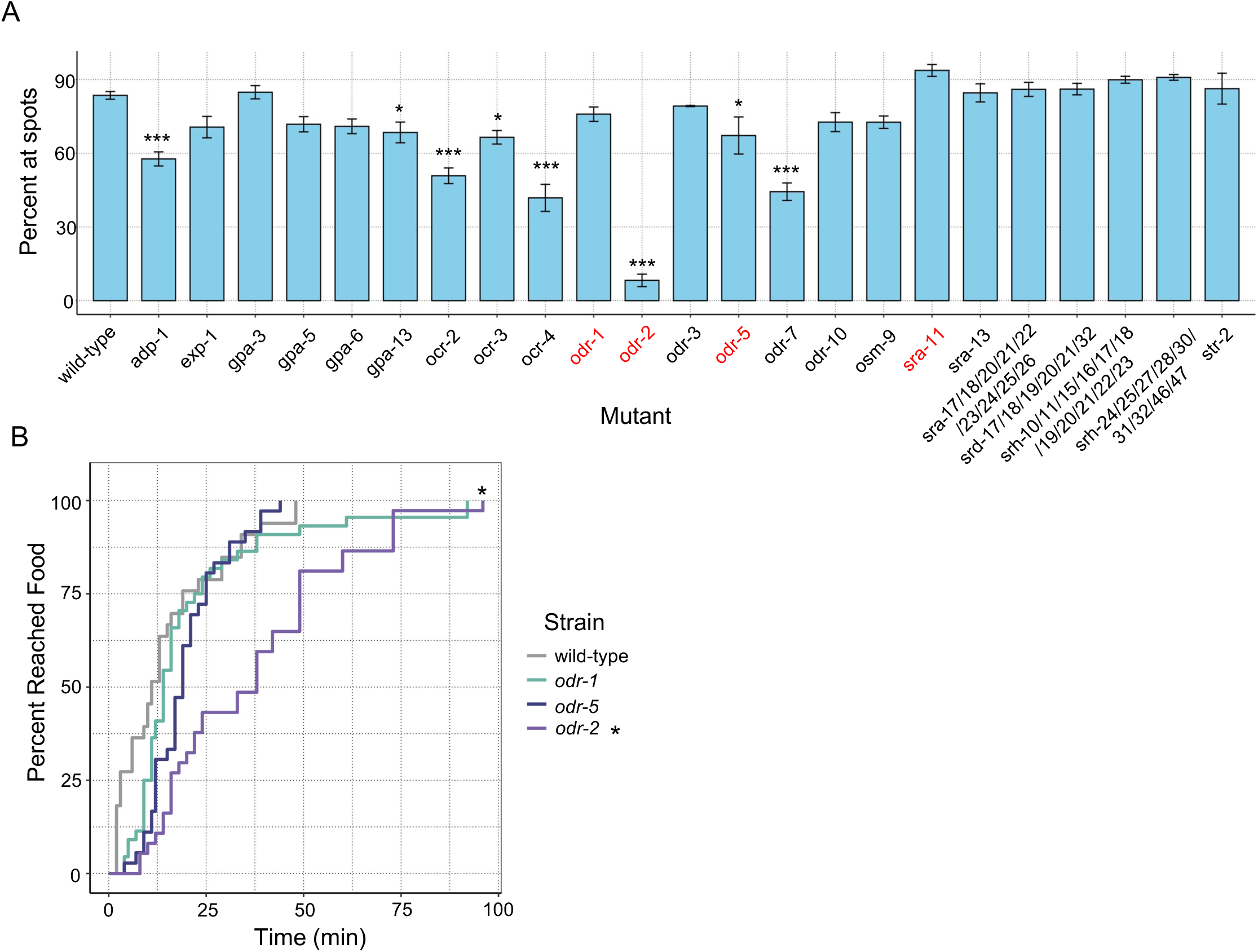
*odr-2* mutants show reduced food chemotaxis. **(A)** Complementary results to those in Fig. 4A, quantifying the percent of worms on either attractant spot. Each column represents 5 technical replicates (plates), each with ∼100-300 worms. *, **, *** indicate *p* < 0.05, *p* < 0.01 and *p* < 0.001, respectively, one-way ANOVA with Dunnett post-hoc. **(B)** Worms were placed near an *E.coli* food source, and the time to reach the source was measured. *, *p* < 0.05, log-rank test.

**Fig. S10.**
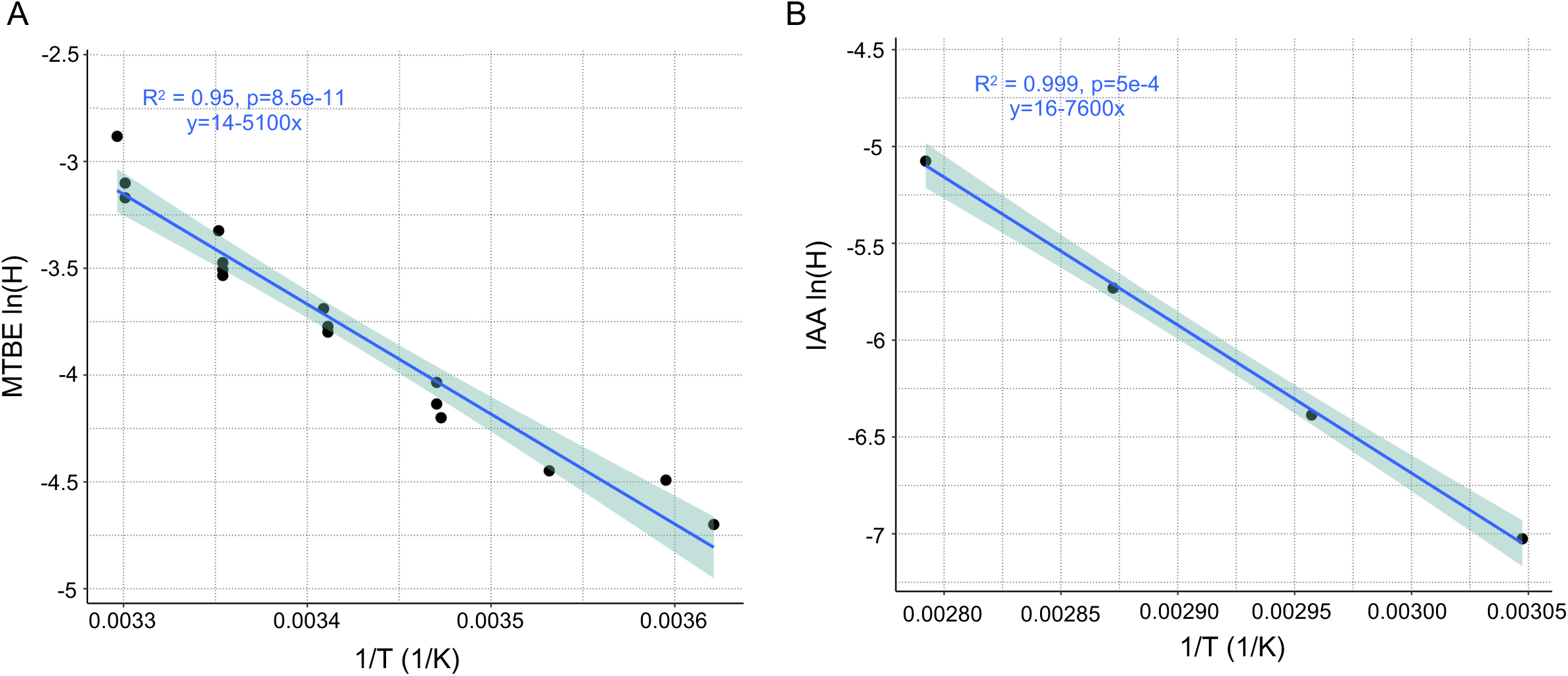
Van’t Hoff plots generated from literature values. **(A)** Curves for tert-butyl methyl ether **(B)** Curves for isoamyl alcohol

## Notes

https://github.com/jpietropaolo579/-Pietropaolo-et-al.-2026-Data-and-Supporting-Files

### Competing Interest Statement

The authors have declared no competing interest.

