## Supplementary Materials for "Attraction to secreted isoamyl alcohol as a signal for beneficial commensals"

**Table S1: Worm Mutants Used in This Study ^a^**

| **Gene(allele)** | **^b^ Effects on IAA Preference (this study)** | | **Previous Reports of Potential Involvement in IAA sensing** | **^c^ Encoded protein (Expression)** |
| --- | --- | --- | --- | --- |
| *adp-1(ky20)* | - | | [1] | - |
| *exp-1(ok1131)* | - | | [2] | GABA-gated cation channel (many neurons) |
| *gpa-3(pk35)* | - | | [3] | Gα (AWA/AWC) |
| *gpa-5(pk376)* | - | | [3] | Gα (many neurons) |
| *gpa-6(pk480)* | - | | [3] | Gα (many neurons) |
| *gpa-13(pk1270)* | - | | [3] | Gα (many neurons) |
| *ocr-2(ak47)* | - | | [4] | TRPV channel (many neurons) |
| *ocr-3(ok1559)* | - | | [5] | TRPV channel (many neurons) |
| *ocr-4(vs137)* | **+** | | [6] | TRPV channel (many neurons) |
| *odr-1(n1936)* | **+++** | | [7, 8] | GC (several neurons, primarily AWB and AWC) |
| *odr-2(n2145)* | **+++** | | [4] | GPI-linked signaling protein (many neurons) |
| *odr-3(n2046)* | - | | [5] | Gα (many neurons) |
| *odr-5(ky9)* | **+++** | | [9] | (AWC) |
| *odr-7(ky4)* | - | | [10] | NHR (AWA) |
| *odr-10(ky225)* | - | | [11] | GPCR (AWA) |
| *osm-9(ky10)* | - | | [1] | TRPV (many neurons) |
| *sra-11(ok630)* | **+++** | | [12] | GPCR (many neurons) |
| *sra-13(zh13)* | - | | [12] | GPCR (many neurons) |
| *str-2(ok3148)* | - | | [2] | GPCR (ASI/AWC) |
| ^d^ ***sra-17(yum1582)*** */18/20/21/22/23/24/25/26* | | - | [13] | **GPCR (ASI/AWA)** + GPCRs expressed in other neurons |
| ^d^ ***srd-17(yum1389)****/18 /19/20/21/32* | | - | [13] | **GPCR (AWC)** + GPCRs expressed in other neurons |
| ^d^ ***srh-10(yum2523)****/11/15 /16/17/18/19/20/21/22/23* | | - | [13] | **GPCR (ASH/ASJ)** + GPCRs expressed in other neurons |
| ^d^ ***srh-28(yum2763) /****24/25/27/30/31/32/46/47* | | - | [13] | **GPCR (ASH/ASJ)** + GPCRs expressed in other neurons |

***^a^*** *Summary of all mutants shown in Figure 4.*

***^b^*** *[IAA] = 0.5 mM*

***^c^*** *Abbreviations: gamma-aminobutyric acid (GABA), G-protein alpha subunit (Gα), transient receptor potential V (TRPV), guanylate cyclase (GC), glycosylphosphatidylinositol (GPI), nuclear hormone receptor (NHR), G-protein coupled receptor (GPCR)*

***^d^*** *Strains with compound disruptions; information provided in rows pertains to gene in boldface.*

**Table S2.** **Genomes Assembly Features**

| **Strain** | **Size (bp)** | **GC%** | **N50/L50** | **Final k-mer Coverage per Contig (median)** | **# Contigs** |
| --- | --- | --- | --- | --- | --- |
| T15 | 4,864,112 | 51.19 | 279980/6 | 1649 | 31 |
| T19 | 4,865,393 | 51.19 | 289462/6 | 1733 | 28 |
| T30 | 4,996,306 | 51.27 | 323452/6 | 1820 | 29 |

**Table S3: Average Nucleotide Identity (ANI) Matrix**

|  | **BIGb0393** | **T14** | **T15** | **T16** | **T19** | **T30** | **V8** |
| --- | --- | --- | --- | --- | --- | --- | --- |
| **BIGb0393** | 100 | 80.4881 | 80.4893 | 82.945 | 80.5039 | 80.5873 | 82.9209 |
| **T14** | 80.5147 | 100 | 99.9952 | 80.5235 | 99.9902 | 99.359 | 80.5338 |
| **T15** | 80.5503 | 99.9911 | 100 | 80.5283 | 99.9989 | 99.3948 | 80.5366 |
| **T16** | 82.9673 | 80.4774 | 80.4675 | 100 | 80.4962 | 80.4399 | 99.9995 |
| **T19** | 80.5985 | 99.9913 | 99.992 | 80.5032 | 100 | 99.4034 | 80.555 |
| **T30** | 80.5771 | 99.3591 | 99.3741 | 80.5058 | 99.368 | 100 | 80.512 |
| **V8** | 82.9288 | 80.4372 | 80.4336 | 99.9995 | 80.4307 | 80.4592 | 100 |

**Table S4: Pantoea strains similarity and differences**

| **Genome A** | **Genome B** | **Avg Identity** | **Aligned Bases** | **Unaligned Bases** | **Total SNPs** | **Total Indels** | **SNPs per Mbp** |
| --- | --- | --- | --- | --- | --- | --- | --- |
| **BIGb0393** | **T14** | 83.37 | 1328112 | 3862186 | 208224 | 10257 | 156781.958 |
| **BIGb0393** | **T15** | 83.37 | 1330491 | 3859807 | 208323 | 10246 | 156576.031 |
| **BIGb0393** | **T16** | 84.32 | 2638728 | 2551570 | 393710 | 19606 | 149204.465 |
| **BIGb0393** | **T19** | 83.39 | 1334215 | 3856083 | 208218 | 10272 | 156060.305 |
| **BIGb0393** | **T30** | 83.41 | 1337644 | 3852654 | 209083 | 10100 | 156306.91 |
| **BIGb0393** | **V8** | 84.31 | 2631830 | 2558468 | 393459 | 19596 | 149500.158 |
| **T14** | **T15** | 99.99 | 4864523 | 1192 | 11 | 1 | 2.261 |
| **T14** | **T16** | 83.45 | 1274408 | 3591307 | 202052 | 9341 | 158545.772 |
| **T14** | **T19** | 99.99 | 4865002 | 713 | 16 | 1 | 3.289 |
| **T14** | **T30** | 99.46 | 4746948 | 118767 | 22910 | 1674 | 4826.259 |
| **T14** | **V8** | 83.44 | 1273381 | 3592334 | 201980 | 9302 | 158617.099 |
| **T15** | **T16** | 83.44 | 1276378 | 3587734 | 202261 | 9352 | 158464.812 |
| **T15** | **T19** | 100 | 4864081 | 31 | 3 | 0 | 0.617 |
| **T15** | **T30** | 99.46 | 4745684 | 118428 | 22901 | 1718 | 4825.648 |
| **T15** | **V8** | 83.44 | 1275347 | 3588765 | 202120 | 9310 | 158482.358 |
| **T16** | **T19** | 83.44 | 1279980 | 5093304 | 202567 | 9394 | 158257.942 |
| **T16** | **T30** | 83.43 | 1274826 | 5098458 | 201775 | 9423 | 158276.502 |
| **T16** | **V8** | 100 | 6371757 | 1527 | 10 | 0 | 1.569 |
| **T19** | **T30** | 99.46 | 4746336 | 119057 | 22939 | 1751 | 4832.991 |
| **T19** | **V8** | 83.43 | 1275458 | 3589935 | 202117 | 9320 | 158466.214 |
| **T30** | **V8** | 83.43 | 1270863 | 3725443 | 201411 | 9348 | 158483.645 |

**^a^** While sequence differences between strains T14 /T15 and between T14 (and T15)/T19 genomes appear small, worms reproducibly preferred T14 slightly over T15 and T14 (and T15) much more strongly than T19 (Fig. 1E), suggesting that they are all distinct.

**Table S5: Oligonucleotides (Spacers) Used for CRISPR-based knock-out**

| **Primer** | **^a^ Sequence (5' → 3')** |
| --- | --- |
| V8-adhE-spacer1-F | ATAACGGCATGGGCATCGTCGAAGATAAAGTGATCAAG |
| V8-adhE-spacer1-R | TTCACTTGATCACTTTATCTTCGACGATGCCCATGCCG |
| V8-adh1-spacer1-F | ATAACTCGTTCGGACGCGCGTGATATCGACTTAGCACG |
| V8-adh1-spacer1-R | TTCACGTGCTAAGTCGATATCACGCGCGTCCGAACGAG |
| V8-adh2-spacer1-F | ATAACTGGAGAACAAGGTTCTCTGATTACACTACAACG |
| V8-adh2-spacer1-R | TTCACGTTGTAGTGTAATCAGAGAACCTTGTTCTCCAG |
| V8-adhT-spacer1-F | ATAACATGAAGGCGTTGGTCAAATCGTGGCGTTAGGTG |
| V8-adhT-spacer1-R | TTCACACCTAACGCCACGATTTGACCAACGCCTTCATG |
| V8-alKJ-spacer1-F | ATAACGCTGGGAACAGCAAGGTTGTGAGGGCTGGGGCG |
| V8-alKJ-spacer1-R | TTCACGCCCCAGCCCTCACAACCTTGCTGTTCCCAGCG |
| V8-ydfG_1-spacer1-F | ATAACTGCAGGCACTGAAAGACGAGCTGGGCGACAATG |
| V8-ydfG_1-spacer1-R | TTCACATTGTCGCCCAGCTCGTCTTTCAGTGCCTGCAG |
| V8-ydfG_2-spacer1-F | ATAACTCGCGCTGGATGTCAGCGATCACCTTGCGGTGG |
| V8-ydfG_2-spacer1-R | TTCACCACCGCAAGGTGATCGCTGACATCCAGCGCGAG |
| V8-ydfG_3-spacer1-F | ATAACGTAAAGAACAGCTGGCGGCGCTGGCGGAACGCG |
| V8-ydfG_3-spacer1-R | TTCACGCGTTCCGCCAGCGCCGCCAGCTGTTCTTTACG |
| V8-ldcC-spacer1-F | ATAACATTCGCAAATGGATGTCAGCATCAACGAAATGG |
| V8-ldcC-spacer1-R | TTCACCATTTCGTTGATGCTGACATCCATTTGCGAATG |
| V8-sgbH-spacer1-F | ATAACATTTTATGTTTCAGCGCGGGCATTGAGGCGGTG |
| V8-sgbH-spacer1-R | TTCACACCGCCTCAATGCCCGCGCTGAAACATAAAATG |
| V8-ilvE_1-spacer1-F | ATAACGTCATCGCGAACACATGCAGCGTCTGCACGACG |
| V8-ilvE_1-spacer1-R | TTCACGTCGTGCAGACGCTGCATGTGTTCGCGATGACG |
| V8-ilvE_2-spacer1-F | ATAACGGATATGTCTGTGGTGATGGTGTATGGGAAGGG |
| V8-ilvE_2-spacer1-R | TTCACCCTTCCCATACACCATCACCACAGACATATCCG |
| V8-tyrB-spacer1-F | ATAACATATCGCTATCTTTAACGGCGCAGGCTTTGAGG |
| V8-tyrB-spacer1-R | TTCACCTCAAAGCCTGCGCCGTTAAAGATAGCGATATG |

**^a^** Red letters denote BsaI-generated sticky ends facilitating ligation into the pBFC0619 vector.

**Table S6: Verification Primers for CRISPR Knockouts**

| **Gene** | **Primer** | **Sequence (5' → 3')** |
| --- | --- | --- |
| ***sgbH*** | Check Fwd | ATGAGCCTTCCTTTGCTCCAGCTGGCGCT |
| *sgbH* | Check Rev | TTAGCACCGGCGTCGAATGCCATGCGG |
| ***ydfG1*** | Check Fwd | ATGATTATTTTTGTTACCGGTGCGACCGCCGGTTTTGGT |
| *ydfG1* | Check Rev | GCATTGTTAACCAGCACATCAATATTGCGCCATTCAGCAGGC |
| ***ldcC*** | Check Fwd | TTGAATATTCTTGCGATCATGGGAGCGCACGGCG |
| *ldcC* | Check Rev | TTCATCGGTGTATTGCCGGATGTGCAGCGCGATATC |
| ***ydfG2*** | Check Fwd | TTGCGGCATATCTGCACTGA |
| *ydfG2* | Check Rev | CCCGCCATAGGACTGACAAA |
| ***adh1*** | Check Fwd | CACGCCCGGCGCTTTGGTTTCTTTTAAAGCGACTTA |
| *adh1* | Check Rev | CAATCCGGTCGGCCACCTGTAACAGCAGGTT |
| ***adhT*** | Check Fwd | CCGTTGGACCGGGCCAGGTGCTGGTTAAAAT |
| *adhT* | Check Rev | AGCTATCAAGACAGTATTCGCAATGGCCGCAAGCTGAGT |
| ***ilvE1*** | Check Fwd | TGGTTAAGTGGGAAGACGCC |
| *ilvE1* | Check Rev | TTGTCTTCAGTTTCGCCGGT |
| ***adh2*** | Check Fwd | TCACAGTTTCCCATCGGCAGATCTGACAGCGTGG |
| *adh2* | Check Rev | CAATCGGCGAGGCATTGATCGGCGAGGTGTTAA |
| ***ilvE2*** | Check Fwd | TTTATGTGAATGGCGAATTTGTCCACCGGGACAACGCT |
| *ilvE2* | Check Rev | ATTTCCTCACGGGTGTGACCAATGTCGAGCTGGATAGA |
| ***ydfG3*** | Check Fwd | CTCCCGCGGTTATAAGGGTC |
| *ydfG3* | Check Rev | TCGCCAGAAACGTTACCGAA |
| ***tyrB*** | Check Fwd | TTCCGGTGCGCTGAAAGTAGGTGCGGATTTCCT |
| *tyrB* | Check Rev | AGCAGCACGATGCTTTGCTTCGGCAGGGTTTT |
| ***alkJ*** | Check Fwd | GCAAGGCGCTGGGTGGTTCTTCATCCATGAACAG |
| *alkJ* | Check Rev | GACCAGTAATTCGCCGTTAAAACCATGATATGCCGGGTCCT |
| ***adhE*** | Check Fwd | AAGTGGGTGCTGAATGGTGT |
| *adhE* | Check Rev | CACCTTTACGCACGATGCTC |

**Table S7: V8 homologs of putative IAA synthesis enzymes**

| **Gene** | **Translated protein** |
| --- | --- |
| *ilvE1* | MSTKKADFIWFNGEMVKWEDAKVSVMSHALHYGTSVFEGVRCYDSHKGPVVFRHREHMQRLHDSAKIYRFPIEASVDELMEACREVLRVNKLKSAYIRPLAFVGDVGLGVNPPDGYTTDVIIAAFPWGAYLGAEALEQGIDAMVSSWNRVAPNTLPTAAKAGGNYLSSLLVGSEARRHGYQEGIALDTNGYISEGAGENLFEVKDGILFTPPFTSSALPGITRDAIIKLAKDMGIEVREQVLSRESLYLADEVFMSGTAAEITPVRSVDGIQVGAGKCGPVTKRIQQAFFGLFTGETEDKWGWLDPVNA |
| *ilvE2* | MSASQSSQAYLQDSRNNNVQVYVNGEFVHRDNATVSIFDSGYVCGDGVWEGLRLVNGKLIALQNHLDRLFAGAASIQLDIGHTREEITAIMFKTLEVNGMTDGAHLRLMITRGRKRTPNQDPRFIIGGATVVCVAEYKVVDYEAKKRGLALFTSSYRTSTPDVFDLRLNSHSRLNLIQALLQALDAGADEALMLDPHGFVASCNSTNFFIVRNGELWTSNGLYCFNGITRQTLIELARQNGLAVYEKPFTLAEALTADEAFVTGTLAGITPVRKLDGRAFDVSQTPVTTQLAQWFQYYLNNI |
| *tyrB* | MFQNVDAYAGDPILSLMETFKQDPRDNKVNLSIGLYYNEQGIIPQLQAVAAAEERLQAAPHQASLYLPMEGFGPYRNAIAPLLFGSDHPMLKAGRIASIQTLGGSGALKVGADFLKRYFPNSNVWVSDPTWENHIAIFNGAGFEVNTYPWYDAETNGVKFDAFIAALKTLPKQSIVLLHPCCHNPTGADLTDAQWDQTVAVLKAQELIPFLDIAYQGFGAGMNEDAYAIRAVAAAGLPALVSNSFSKIFSLYGERVGGLSIVCDSAEEAGRVLGQLKATVRRNYSSPPNFGAQVVSCVLNDEALLNNWLAEVEAMRLRIIEMRQALVAVLREKLPGQNFDYLLKQRGMFSYTGLSAQQVDRLRDEFGVYLIASGRMCVAGLNSRNVHQVAEAFAAVM |
| *ldcC* | MNILAIMGAHGVFYKDEPIRELDAALKLQGFQTVYPTNASDLLKLIEHNPRICGVIFDWDDYSLELCSEINQLNEYLPLYAFINTHSQMDVSINEMRMALHFFEYALSAADDIALHIRQYTDEYIDKITPPLTKALFTYVKEGKYTFCTPGHMGGTAFQKSPVGSLFYDFFGANTLKADISISVTELGSLLDHTGPHLEAEEYVARTFGAEQSYMVTNGTSTSNKIVGMYAAPAGSTVLIDRNCHKSLTHLLMMSDIIPLWLKPTRNALGILGGIPQREFTRDSIQHKVDVTPRASWPVHAVITNSTYDGLLYNTQYIKQTLDVPSIHFDSAWVPYTNFHPIYAGKSGMSGDRVPGKVFYETQSTHKLLAAFSQASLIHIKGDYDEQTFNEAYMMHTTTSPNYAIVSSIETAAAMLRGNPGKRLINRSVERALHFRREIQRLREESDGWFFDIWQPEQVDEAECWPIQPGEEEWHGFTQADRDHMYLDPIKVTILTPGMSELGVMAEEGIPAALVAKFLDQRGVVVEKTGPYNLLFLFSIGIDKTKAMSVLRGLTEFKRAYDLNLRVKNMLPDLYAEDPDFYRNMRIQTLAQGIHQLILQHDLPRLMLKAFDVLPEMKMTPHQAFQQQVKGQVETVDIRELVGRISANMILPYPPGVPLVMPGEMITEQSRAVLDFLLMLCTIGRHYPGFETDIHGATLTEDGRYLVRVLKGENDH |
| *sgbH* | MSLPLLQLALDHTDLDAALSTAQQLHQQVDIIEAGTILCFSAGIEAVRRLRQQHPGKTLVADFKVADAGATLARMAFDAGANWMTVICAAPLATFATALEVAREYQGDIQIELFGHWTLEDARQWRSLGLTQAIYHRGRDAQASGQQWSQQDLDAMQALSDMGFALSITGGITPAELPQFRNIDVKAFIAGRALSDSVTGIETASQFHQAIANIWGTP |
| *ydfG1* | MIIFVTGATAGFGQSITRRFIATGHKVIASGRRAERLQALKDELGDNLYTVQLDVRNRAAIEEAIAALPAEWRNIDVLVNNAGLALGVEPAHKANIEDWENMIDTNNKGLVYMTRALLPAMVERNVGHIINIGSIAGSWPYLGGNVYGATKAFVRQFSLNLRTDLHGTALRVTDIEPGLVGGTEFSNVRFKGDDGKADKVYEGTTALTAEDVTEAVYWVATLPKHVNINTLEMMPVTQTLAGLKVHKE |
| *ydfG2* | MLIFITGATAGFGWDLALRYAQHGHRVIATGRRQDKLDQLKAAGGENIFTLALDVSDHLAVEGLLEQLPLAWRDIDVLINNAGLALGLEPAQKASIDDWNTMIDVNIKGLVHVTRVLLPGMVERNRGHIINLGSTAGSWPYAGGNVYGASKAFVRQFSLNLRTDLFGTAVRVTNIEPGLVGGTEFSSVRFKGDENRVEQTYADTQPLTPADITEAIWWVSNLPAHVNINTLEMMPVCQSYGGLRVAKQG |
| *ydfG3* | MPFSDYKTALVTGASAGMGEAIVERLCQEGITVHAVARRKEQLAALAERTGCIPHAVDVSDLSALTALCQDLQIDILVNNAGLSHPGSILDADENVIESQVDVNLRAVLHLCRLLVPGMVARDRGHVFNITSIAAIYNFGGNSVYHATKAGVHALSRQLRVDCYGKRVRITEICPGRVATDIFGNVSGDHEDARRRFIDGFELPQAKDIADCVAFALAAPVAVNIGNIEITPTLQVPGGLSTMRPGDRTGS |
| *adhE* | MAVTNVAELNALVERVKKAQREYANFSQEQVDAIFRAAALAAADARIPLAKMAVAESGMGIVEDKVIKNHFASEYIYNAYKDEKTCGVLDTDDTFGTITIAEPIGLICGIVPTTNPTSTAIFKALISLKTRNGIIFSPHPRAKDATNKAADIVLQAAIAAGAPKDIIGWIDAPSVELSNQLMHHPDINLILATGGPGMVKAAYSSGKPAIGVGAGNTPVVVDETADIKRAVASILMSKTFDNGVICASEQSVIVVDSAYDAVRERFATHGGYMLQGKELKAVQDIILKNGALNAAIVGQPAPKIAEMAGITVPANTKILIGEVKLVDESEPFAHEKLSPTLAMYRAKDFQDAVDKAEKLVAMGGIGHTSCLYTDQDNQNERVHYFGDKMKTARILINTPASQGGIGDLYNFKLAPSLTLGCGSWGGNSISENVGPKHLINKKTVAKRAENMLWHKLPKSIYFRRGSLPIALEEVATDGAKRAFIVTDRFLFNNGYADQVTRVLKSHGIETEVFFEVEADPTLSIVRKGAEQMNSFKPDVIIALGGGSPMDAAKIMWVMYEHPETHFEELALRFMDIRKRIYKFPKMGVKARMIAITTTSGTGSEVTPFAVVTDDATGQKYPLADYALTPDMAIVDANLVMDMPRSLCAFGGLDAVTHSLEAYVSVLANEYSDGQALQALKLLKENLPASYKEGAKNPVARERVHNAATIAGIAFANAFLGVCHSMAHKLGSEFHIPHGLANALLICNVIRYNANDNPTKQTAFSQYDRPQARRRYAEVADHLGLSAEGDRTAQKIEKLLAWLEEMKTQLGIPTSIREAGVQEADFLAKVDKLADDAFDDQCTGANPRYPLIAELKQIMLDTFYGREFVETTAEVTEAVDLQPVKSVKTEKKTKKA |
| *adh1* | MRYAHPGTPGALVSFKATYGNYIAGKFTEPLSGQYFTNTSPVDGSDIAQFPRSDARDIDLALDAAHQAADAWGKTSAQHRANLLLQVADRIEANLEMLAVAESWDNGKPIRETLNADLPLAADHFRYFAGCLRAQEGSTAEIDEHTVAYHFHEPLGVVGQIIPWNFPLLMAAWKLAPALAAGNCVVLKPAEQTPLGITLLLELIGDLFPAGVLNVVQGFGREAGEALATSKRIAKIAFTGSTPVGRHIMACAAENIIPCTVELGGKSPNIYFADVMDGEPEFIEKAVEGLVLGFFNQGEVCTCPSRALIHESIYAPFMERVMAKIATIRRGDPLDTETMIGAQASRQQFDKILSYIDIARQEGGQILTGGERASITPALDNGFYIQPTLIKGDNQMRCFQEEIFGPVIGVTTFKDEAEALEIANQTQFGLGAGVWTRNTNLAYRMGRSIKAGRVWTNCYHIYPAHAAFGGYKQSGVGRETHKMALNAYQQTKNLLVSYDIAPLGLF |
| *adh2* | MRYAAPGEQGSLITLQQRYGNFINGEFVPPVNGNYFVNTSPINASPIGEFPRSDRDDVDNAIAAAHKAADAWGKTSPQARSLVLLKIADRLEENLEYLAVNETWDNGKPVRETLAADMPLAVDHFRYFAGCVRAQEGTAAEIDEFTASYHFHEPLGVVGQIIPWNFPLLMAAWKLAPALGAGNCVVLKPAEQTPLSITLFMELIKDLVPPGVINVVHGYGKEAGEALASHPGIAKIAFTGSTATGGHILELAAKSLIPSTVELGGKSPNIFFEDIMQAEESFIEKAAEGVVLGFLNQGEVCTCPSRALVQESIFEPFMAAVMKRIKTIKRGNPLDTETMIGAQASQQQFDKILSYLEIAQQEGAQVLHGGGIESVGEEFASGYYIQPTLLKGSNSMRVFQEEIFGPVVGVLTFKDEAEAIALANDSMYGLGAGVWTRDINRAYRMGRAIKAGRVWTNCYHLYPAHAAFGGYKKSGIGRETHKMMLDHYQQTKNLLVSYSTQPLGFF |
| *adhT* | MTMKIKSTMKAAVVKSFGEPLVIEDVPVPSVGPGQVLVKIAATGVCHTDLHAAEGDWPIKPNPPFIPGHEGVGQIVALGEGVKHLKMGDRVGVPWLYSACGHCEYCLDSWETLCLSQQNAGYSVNGSFAEYCLADANYVGILPDNIEFNEIAPILCAGVTVYKGLKMTDTKPGDWVVISGIGGLGHMAVQYAVAMGLNVAAVDIDDEKLAFAQRLGASVVANAKNVDPAKVFHESFGGAHGVLVTAVSPKAFEQALGTMRRGGTMVLNGLPPGKFDLSIFDMVLDGITVRGSIVGTRKDLQEALDFAGRHKVKANVAVEPLVNINDIFARMHAGKIEGRIVVDMSL |
| *alkJ* | MNNSNTYDYIIIGGGSAGSVLAARLAEQADLKICLIEAGSRDETPRIQTPAGTITLYKSKKFSWNFYSTPQKSLGGRQLHVPRGKALGGSSSMNSMIYIRGLPSDYDRWEQQGCEGWGWNNVLPWFKRSEKNLLSQDPAYHGFNGELLVDKPRDPNPVSALFVAAGKRVGLAENTDFNGKSLAGVGIYNVTQKDGKRLSSYRAFLHPHIGQSNLTVMTDCTVQTLIIEDKVVKGVRITEHGRDQPTSILCRREVILSAGSIGSPHILLKSGIGPAAELEAAGIPLMHPLPGVGKNLQDHLDGLVTVRSGNPLTLGFSLAAWKPILTSPFNYLFRRKGWLTTNYVEAGGFAATKLSSDEPDIQFHFVPGYRSHRGRLFEWGHGYAIHTCVLRPKSIGALQLTRDGQIAIDFNFLADPYDASVLVEGIKVARNILAQPEFAALRGEEMLPGKHIQTDEQLHQYVKEYCATVFHPVGTCKMGRDEMSVVAPDTLKVYGVENLRVADASIMPSLISGNTNAPSIMIGERAASMILQGTPVADTETNKENAYA |

**Quantification of IAA and 2-PE Concentrations from Headspace GC-MS Peak Areas**

**(A) Mole Balance of MTBE in the Sample Apparatus**

$$\left( Mole Balance \right) Equation 1.1:n_{total}= n_{liq}+n_{gas}$$

n = moles of analyte

A concentration times the volume of the sample is equal to the total moles in a sample, which leads to equation 1.2:

$$Equation 1.2:n_{total}= C_{liq,T}V_{liq,T}+C_{gas,T} V_{gas,T}$$

C = concentration of analyte

V = volume of phase

$$\left( \mathrm{Henr}y^{'}s Law \right) Equation 2.1:C_{gas,T}=C_{liq,T}\times H_{T}$$

H = Henry’s Law Constant (dimensionless)

Plugging Equation 2.1 into equation 1.2 yields equation 1.3

$$Equation 1.3:n_{total}= C_{liq,T}V_{liq,T}+C_{liq,T}H_{T}V_{gas,T}$$

Rearranging equation 1.3 and plugging it into Henry’s Law results in equation 2.2

$$Equation 2.2:\frac{n_{total}H_{T}}{V_{liq,T}+H_{T} V_{gas,T}}=C_{gas,T}$$

The only missing value now is $H_{T}$. Once we find that, we can plug in the values to find the vapor pressure of MTBE and by extension the analytes of interest.

**(B) Calculation of MTBE’s Henry’s Law Constant (at 55ºC) from Literature Values**

Using dimensionless Henry’s law constants from previous papers [14, 15], a Van’t Hoff plot (Fig. S10A) allowed extrapolation of Henry’s constant for MTBE from published values for different temperature to a **Henry constant for MTBE at 55ºC of 0.0156, and at 20ºC of 0.0241.**

**(C) Calculation of Relative Response Factors (RRFs)**

In order to calculate headspace concentrations of analytes from the internal standard MTBE, relative response factors should be incorporated describing differences in ionization efficiencies, matrix effects, injection effects, etc.. RRFs for the two analytes, were calculated from the liquid calibration, yielding $\boldsymbol{RRF}_{\boldsymbol{IAA}}\boldsymbol{=2.90,}\boldsymbol{RRF}_{\boldsymbol{2-PE}}\boldsymbol{=0.0886}$.

$$Equation 3.1:\frac{A_{a}}{A_{IS}}=RRF\frac{C_{a}}{C_{IS}}$$

$A_{a}$= Peak area of analyte

$A_{IS}$= Peak area of internal standard

RRF = Relative Response Factor

$C_{a}$= Concentration of analyte

$C_{IS}$= Concentration of internal sample

In constructing calibration curves for liquid quantification, we plotted $\frac{A_{a}}{A_{IS}} vs. C_{a}$ (Fig. S3). This means that the slope of the regression is $\frac{C_{IS}}{\mathrm{RRF}}$, which implies equation 4.

$$Equation 3.2: RRF=\frac{C_{IS}}{\mathrm{slope}}$$

**(D) Quantitating Analytes at 55ºC**

Plugging in known values for equation 2.2 we can solve for the concentration of MTBE in the headspace of the samples:

$$Equation 2.2:\frac{n_{total}H_{55}}{V_{liq,55}+H_{55} V_{gas,55}}=C_{gas,55}= \frac{16.79mM*20mL*0.156}{20mL+0.156*30mL}=2.13 mM$$

Now plugging in known values into equation 3.1 yields the headspace concentrations of analytes at 55ºC sampling conditions. As an example calculation using IAA with a V8 headspace sample:

$$Equation 3.1:\frac{A_{a}}{A_{IS}}=\frac{A_{IAA}}{A_{MTBE}}={RRF}_{IAA}\frac{C_{IAA,55}}{C_{MTBE,55}}\to\frac{3.98e7}{7.67e9}=2.90\frac{C_{IAA,55}}{2.13mM}\to\boldsymbol{C}_{\boldsymbol{IAA gas,55ºC}}\boldsymbol{=3.81}\boldsymbol{e-3 mM}$$

**(E) Calculating IAA and 2-PE Henry’s law Constant Calculations**

Using dimensionless Henry’s law constants from a previous paper [16] a Van’t Hoff plot was constructed for IAA (Fig. S10B), yielding $\mathbf{H}_{\mathbf{IAA, 55ºC}}\boldsymbol{=8.69}\boldsymbol{e-4,}\mathbf{H}_{\mathbf{IAA,20ºC}}\boldsymbol{=5.39}\boldsymbol{e-5}$.

Published data for dimensionless Henry’s law constants for 2-phenylethanol is less available, which required using the Vant-Hoff equation with the $\frac{{\Delta H}_{\mathrm{sol}}}{R}$parameter, as described in [17], and subsequently converting to a dimensionless version yielding:

$$H_{PE,55ºC}= \boldsymbol{2.29}\boldsymbol{e-6}, H_{PE,20ºC}=\boldsymbol{3.52}\boldsymbol{e-6}$$

**(F) Quantitating Analytes at 20ºC**

Using equation 2.2 **(A)** in reverse we can calculate$n_{total}$ for IAA in the tube based on its concentration at 55ºC at the gas phase (and continuing with the example in **(D)**, yielding:

$$Equation 4:C_{gas,T}\left( \frac{V_{liq,T}}{H_{T}}+V_{gas,T} \right)=n_{total}=3.81e-3mM\left( \frac{20mL}{8.69e-4}+30mL \right)=8.80 e-5 moles$$

and with the calculated Henry’s law constant find it concentration in the headspace:

$$Equation 2.2:\frac{8.80 e-5 mol* 5.39e-5}{20mL+(5.39e-5*30mL)}=\boldsymbol{C}_{\boldsymbol{IAA gas,20ºC}}\boldsymbol{=2.37}\boldsymbol{e-4}\boldsymbol{mM}$$

**(G) Key Assumptions**

1. The sampling apparatus is a closed system.

a. The sampling apparatus consisted of a sealed 50 mL Falcon tube. A small puncture was made in the tube cap using a needle of the same gauge as the SPME fiber to allow insertion. The fit between the fiber and the puncture was extremely tight, and removal of the fiber resulted in a brief release of pressurized steam, indicating negligible leakage during incubation and sampling.

2. Gas–liquid equilibrium is achieved prior to sampling.

a. Samples were incubated at the measurement temperature for 20min to allow gas–liquid equilibrium to be established before SPME exposure. Henry’s law and the associated mass-balance relations are therefore applicable.

3. The system remains closed with respect to analyte mass.

a. The total number of moles of analyte ($n_{total}$) is conserved within the sampling vessel. No significant loss of analyte due to leakage, adsorption, degradation, or biological consumption/production occurred during incubation or sampling.

4. SPME sampling does not significantly perturb headspace concentrations.

a. SPME fibers are extremely tiny. Therefore, it is assumed that analyte uptake by the fiber did not measurably deplete the headspace, and GC–MS response remained proportional to headspace concentration under the sampling conditions used.

5. Henry’s law constants for air–water systems approximate those for air–conditioned media.

a. Henry’s law constants reported for air–water systems were used as approximations for air–bacterial conditioned media. While salts and metabolites present in the media may slightly alter activity coefficients, these effects are assumed to be small relative to the observed concentration ranges and acceptable for the purposes of this study.

6. Temperature dependence of Henry’s law constants follows Van’t Hoff behavior.

a. Henry’s law constants at temperatures other than the reference temperature were calculated using a Van’t Hoff relationship and published temperature-dependent coefficients. This approach is widely accepted for dilute systems; any deviation from ideal behavior is expected to be minor.

7. Ideal gas behavior applies in Henry’s Law Constant unit conversions.

a. Conversion between pressure-based and concentration-based Henry’s law constants assumes ideal gas behavior. Given the low analyte partial pressures, near-atmospheric total pressure, moderate temperatures, and a pure air/water system in [17], deviations from ideality are negligible.

8. Liquid and gas volumes are constant during sampling.

a. Liquid and headspace volumes were treated as constant for each calculation. Thermal expansion of phases and volume displacement by the SPME fiber were negligible relative to total system volume.

9. The system is isothermal during sampling.

a. The liquid and headspace phases were assumed to be at uniform temperature during sampling, and the measured incubation temperature was taken to represent both phases. This is reasonable as the sample was maintained in a constant temperature water bath.
